# Role of Dual Specificity Phosphatase 1 (DUSP1) in influencing inflammatory pathways in macrophages modulated by *Borrelia burgdorferi* lipoproteins

**DOI:** 10.1101/2024.11.20.624562

**Authors:** Venkatesh Kumaresan, Chiung-Yu Hung, Brian P. Hermann, J. Seshu

**Affiliations:** Department of Molecular Microbiology and Immunology, South Texas Center for Emerging Infectious Diseases, The University of Texas at San Antonio San Antonio, TX-78249; Department of Neuroscience, Developmental and Regenerative Biology, The University of Texas at San Antonio San Antonio, TX-78249

**Keywords:** Lyme disease, Lipoprotein, bone marrow-derived macrophages, scRNA-Seq, Dual-Specificity Phosphatases, mitochondrial oxidative stress

## Abstract

*Borrelia burgdorferi (Bb)*, the spirochetal agent of Lyme disease, has a large array of lipoproteins that play a significant role in mediating host-pathogen interactions within ticks and vertebrates. Although there is substantial information on the effects of *B. burgdorferi* lipoproteins (*Bb*LP) on immune modulatory pathways, the application of multi-omics methodologies to decode the transcriptional and proteomic patterns associated with host cell responses induced by lipoproteins in murine bone marrow-derived macrophages (BMDMs) has identified additional effectors and pathways. <u>S</u>ingle-<u>c</u>ell <u>RNA-Seq</u> (scRNA-Seq) performed on BMDMs treated with various concentrations of borrelial lipoproteins revealed macrophage subsets within the BMDMs. Differential expression analysis showed that genes encoding various receptors, type I IFN-stimulated genes, signaling chemokines, and mitochondrial genes are altered in BMDMs in response to lipoproteins. Unbiased proteomics analysis of lysates of BMDMs treated with lipoproteins corroborated several of these findings. Notably, <u>du</u>al <u>s</u>pecificity <u>p</u>hosphatase <u>1</u> (*Dusp1*) gene was upregulated during the early stages of BMDM exposure to *Bb*LP. Pre-treatment with benzylidene-3-cyclohexylamino-1-indanone hydrochloride (BCI), an inhibitor of both DUSP1 and 6 prior to exposure to *Bb*LP, demonstrated that DUSP1 negatively regulates NLRP3-mediated pro-inflammatory signaling and positively regulates the expression of interferon-stimulated genes and those encoding *Ccl5*, *Il1b*, and *Cd274*. Moreover, DUSP1, IkB kinase complex and MyD88 also modulate mitochondrial changes in BMDMs treated with borrelial lipoproteins. These findings advance the potential for exploiting DUSP1 as a therapeutic target to regulate host responses in reservoir hosts to limit survival of *B. burgdorferi* during its infectious cycle between ticks and mammalian hosts.

**Importance:** *Borrelia burgdorferi*, the agent of Lyme disease, encodes numerous lipoproteins that play a crucial role as a pathogen associated molecular pattern affecting interactions with tick- and vertebrate-host cells. Single cell transcriptomics validated using unbiased proteomics and conventional molecular biology approaches have demonstrated significant differences in gene expression patterns in a dose- and time-dependent manner following treatment of murine bone marrow derived macrophages with borrelial lipoproteins. Distinct populations of macrophages, alterations in immune signaling pathways, cellular energy production and mitochondrial responses were identified and validated using primary murine macrophages and human reporter cell lines. Notably, the role of Dual Specificity Phosphatase 1 (DUSP1) in influencing several inflammatory, metabolic and mitochondrial responses of macrophages were observed in these studies using known pharmacological inhibitors. Significant outcomes include novel strategies to interfere with immunomodulatory and survival capabilities of *B. burgdorferi* in reservoir hosts affecting its natural infectious life cycle between ticks and vertebrate hosts.

## Introduction

Lyme disease is the most common tick-borne infectious disease in the US with 63,000 cases reported to Centers for Disease Control and Prevention (CDC, Atlanta) with around 476,000 cases estimated to occur each year in the US [1]. The spirochetal agent of Lyme disease, *Borrelia burgdorferi (Bb),* is transmitted to humans following the bite of infected *Ixodes scapularis* ticks [2]. Lyme disease is characterized by an inflammatory skin lesion called erythema migrans at the site of tick bite and the bacterium disseminates to deeper tissues leading to Lyme arthritis, carditis and neuroborreliosis [3–5]. While most cases of Lyme disease are cured with a course of oral antibiotics, a minority of patients in spite of antimicrobial therapy continue to report a range of non-specific symptoms referred to as Post-Treatment Lyme Disease Syndrome (PTLDS) [6–8]. The molecular basis of PTLDS is unclear although the nature of host response to *Bb* infection, co-infection with other tick-borne pathogens and persistence of borrelial antigens such as its peptidoglycan or lipoproteins are likely to contribute to clinical manifestation of PTLDS [9–12].

*Bb* has several unique cellular features as a prokaryotic pathogen such as an outer membrane composed of numerous surface exposed lipoproteins [13–16]; glycolipids, absence of lipopolysaccharide [17–19]; a unique peptidoglycan cell wall [20]; and components incorporating host-derived cholesterol and lipids [3, 21] in addition to a variety of periplasmic, membrane bound and soluble proteins mediating interactions with mammalian [22, 23] and tick host cells [24]. Many of these pathogen-derived determinants play a key role in the transmission, colonization, pathogenicity and survival of *Bb* within a wide range of highly divergent hosts [25]. Several past studies on interactions of *Bb* with cells from mammalian and tick hosts have provided a trove of knowledge on the pathogen-specific gene and protein expression profiles and the regulatory mechanisms that facilitate the survival of the spirochetes in tick and mammalian hosts [26, 27]. Lipoproteins, which make up 8% of the *B. burgdorferi* genome, play a key role in host-pathogen interactions. In *B. burgdorferi* strain B31, around 120 potential lipoproteins are encoded, with 52 on the outer surface and 23 in the periplasm, driving the inflammatory and pathogenic effects in infected mammalian hosts. [28]. Notably, several major surface exposed lipoproteins such as <u>O</u>uter <u>s</u>urface <u>p</u>rotein <u>A</u> (OspA) [29, 30], <u>O</u>uter <u>s</u>urface <u>p</u>rotein <u>C</u> (OspC)[29], fibronectin binding proteins such as BBK32, RevA [31, 32] and decorin binding proteins A and B (DbpA/B)[33], among others, mediate interactions with host matrices and host cell surfaces to facilitate colonization and eventual dissemination of *Bb* within an infected host [26, 31, 34]. Plasticity of lipoproteome of *Bb* also contributes to pathogen fitness within 1) a wide range of hosts, 2) microenvironments within each host and 3) capability to link metabolic constraints to pathogenic attributes. Hence, targeting host cell signaling pathways engaged by borrelial lipoproteins are likely to provide novel targets to limit pathogen burden or counteract specific pathogenic outcomes in infected hosts.

The surface exposed lipoproteins serve as the predominant Pathogen Associated Molecular Patterns (PAMPs) driving the innate and subsequent adaptive immune responses of the mammalian hosts via interactions with <u>T</u>oll-like receptors (TLRs) and non-TLR receptors on host cell surfaces [35, 36]. Among the 13 known TLRs (10 in human and 12 in mice), borrelial components interact with TLR1/TLR2 (diacylated/triacylated lipoproteins), TLR4 (glycolipids), TLR5 (flagellin), TLR6 (diacylated/triacylated lipoproteins), TLR7 (ssRNA), TLR8 (bacterial RNA) and TLR9 (nucleic acids in endosomes of macrophages and monocytes)[37–39]. The fatty acid chains in the N-terminus of *Bb*LP that serve as the TLR2/1 PAMP are tethered in the outer membrane, which results in reduced recognition of lipoproteins in intact spirochetes at the host cell surface [13, 40]. However, upon phagocytosis and degradation of spirochetes in the phagosomes of macrophages, the *Bb*LP and borrelial nucleic acids are readily recognized by TLRs or intracellular sensors [41–43]. All TLRs, except TLR 3 (dsRNA), are known to signal via MyD88 [44]. Further, MyD88 interacts with IRAKs and TRAFs to activate the Ubc13/TAK1 pathway. TAK1 complex activates IKK kinase complex comprising of IKKα and IKKβ and a regulatory component NEMO/IKKγ to phosphorylate IkB, which is degraded allowing translocation of NF-κB to the nucleus for transcriptional regulation of genes encoding inflammatory effectors [39, 45].

Pathways independent of MyD88 (via TLR3) or NF-κB (via TLR7//TLR8/TLR9/MyD88 dependent) are also involved in the expression of Type1 IFN genes/IFN stimulated genes (ISGs). Moreover, expression of ISGs has been shown to be independent of two critical adaptor proteins of TLRs namely MyD88 and TRIF[46–48]. Recently, it has been shown that IFN-1 is induced following co-localization of internalized spirochetes with cyclic GMP-AMP synthase (cGAS) in mouse macrophages and is dependent on Stimulator of Interferon Genes (STING) - a sensor of intracellular DNA [43]. It remains to be determined if host cell derived (mitochondrial or nuclear) or *Bb*-derived nucleic acids drive the cGAS/STING activation leading to ISG expression in murine macrophages.

Macrophages are one of the predominant innate immune cells in the mammalian host, playing a vital role in the early recognition of pathogens. They also significantly influence tissue-specific responses to infection, affecting pathogen colonization and immunomodulatory events across multiple organs [49]. Specifically, during *Bb* infection, macrophages contribute to Lyme arthritis [50], Lyme carditis [51] and neuroborreliosis [5, 52]. Furthermore, the phagocytosis of an extracellular pathogen such as *Bb* modulates the immune responses, contributing to the complexity of pathogenesis and survival of *Bb* [41]. Hence, interrogation of effects of borrelial lipoproteins using murine bone marrow derived macrophages at the single cell level are likely to provide additional host cell factors that influence cytokine, chemokine and inflammatory responses that either contribute to survival of pathogen within reservoir hosts or be cleared from accidental hosts. Single-cell RNA-Seq (scRNA-Seq) analysis enables the study of transcriptional changes in small populations of cells at single cell resolution; identification of distinct cell subsets, heterogeneity within defined cell populations, and uncovering developmental branch points or unknown pathways in response to a variety of signals [53–56]. Interactions between macrophages and fibroblasts in the ankle joints of mice infected with *B. burgdorferi* as well as differential gene expression patterns influencing inflammatory and non-inflammatory processes in various cell populations were recently observed using scRNA-Seq methodology [57]. Moreover, scRNA-Seq analysis combined with B cell receptor sequencing revealed an increased number of B cells in human skin lesions with erythema migrans, providing immunophenotyping of inflammatory responses at the site of tick bites and deposition of Lyme spirochetes [58].

Prior studies from our group using *ex vivo* infection of splenocytes with *Bb* revealed a role for the Dual Specificity Phosphatase 1 (DUSP1) in the Caspase-3 dependent apoptosis of bone marrow derived neutrophils using scRNA-Seq analysis [59]. Extending these studies, we determined the effects of purified borrelial lipoproteins (*Bb*LP) in modulating proinflammatory cytokine levels, altering mitochondrial responses and cell-cell interactions of murine bone marrow derived macrophages using scRNA-Seq analysis and subsequently validating the differentially expressed genes and their regulatory pathways using conventional tools and methodologies. Upon *Bb*LP treatment in BMDMs, DUSP1 was upregulated within one hour of treatment, followed by a rapid decline at both the mRNA and protein levels. The significance of DUSP1 was further confirmed using a known pharmacological inhibitor of DUSP1/6 in borrelial lipoprotein-mediated responses, linking DUSP1 levels to PD-L1, CCL5, IL-1B, and the NLRP3-mediated induction of CXCL1 and CXCL2. In addition, DUSP1 mediated effects on ISG transcripts via perturbations in the host cell mitochondria/mitochondrial DNA adds to the diversity of early innate responses. These studies identify additional targets for regulating the innate immune responses of mammalian hosts, which are likely to provide avenues to interfere with pathogen survival and colonization in reservoir hosts that sustain *B. burgdorferi* in its natural infectious cycle.

## Results

### Comparative scRNA-Seq analysis of BMDMs treated with *Bb*LP

To analyze molecular changes in macrophages at a single cell level, borrelial lipoproteins (*Bb*LP) were purified from *B. burgdorferi* strain B31 employing detergent partitioning method using Triton X-114 (**Supp-Fig. 1**) followed by treating murine BMDMs with 1 µg/mL of *Bb*LP for 1 (LP1) and 4 hours (LP4) with untreated cells maintained as control (Ctrl). Approximately, 5000 cells from each sample were subjected to scRNA-Seq to determine the cellular and transcriptional profile in *Bb*LP treated BMDMs. Cell Ranger analysis was conducted on the FASTQ files, and the data visualization and analyses were performed using the Seurat packages and LOUPE Browser software. The count analysis identified 6,125, 5,119, and 5,039 barcodes for Ctrl, LP1, and LP4, respectively. Quality control steps were implemented to exclude cells with a high percentage of mitochondrial genes (>5%). Further quality control steps were applied to retain cells with UMI counts > 500, genes > 500, and mitochondrial UMIs (mouse) < 5%, ultimately including 12,111 cells for downstream analysis.

All three samples (Ctrl, LP1, and LP4) were integrated, and UMAPs were generated using Harmony-based clustering in Seurat, identifying eleven distinct clusters (**Fig. 1a**). Differential gene expression (DEG) analysis revealed that clusters 1, 3, 4, 7, and 9 shared genes related to phagocytosis, while clusters 0, 2, and 6 were enriched for genes involved in antigen presentation. Clusters 5, 8, and 10 were associated with cell cycle-related genes, distinct from other clusters (**Fig. 1b**). A bubble plot of selected genes showed that phagocytosis-related genes, such as *Apoe* and *Slc9a9*, were predominantly expressed in phagocytosis-associated clusters, while *Fgf13* and *Ccnd1* were enriched in antigen presentation clusters. In contrast, cell cycle-related genes, including *Hist1h1b* and *Hist1h2ap*, were uniquely expressed in clusters associated with cell cycle regulation (**Fig. 1c**). A heatmap comparing the top DEGs between phagocytosis, antigen-presentation, and cell-cycle-related macrophages was generated by grouping similar clusters. (**Fig. 1d**). The UMAP feature plots of these hub genes demonstrated distinct expression patterns for each macrophage subset (**Fig. 1e**), indicating that BMDMs exhibit functionally specialized macrophage subpopulations, with a high degree of diversity in their roles. In addition, we performed differential expression analysis using LOUPE browser and observed comparable subsets of BMDMs as shown in the results presented in the supplementary section (**Supp-Fig. 2 and 3**).

**Figure 1:**
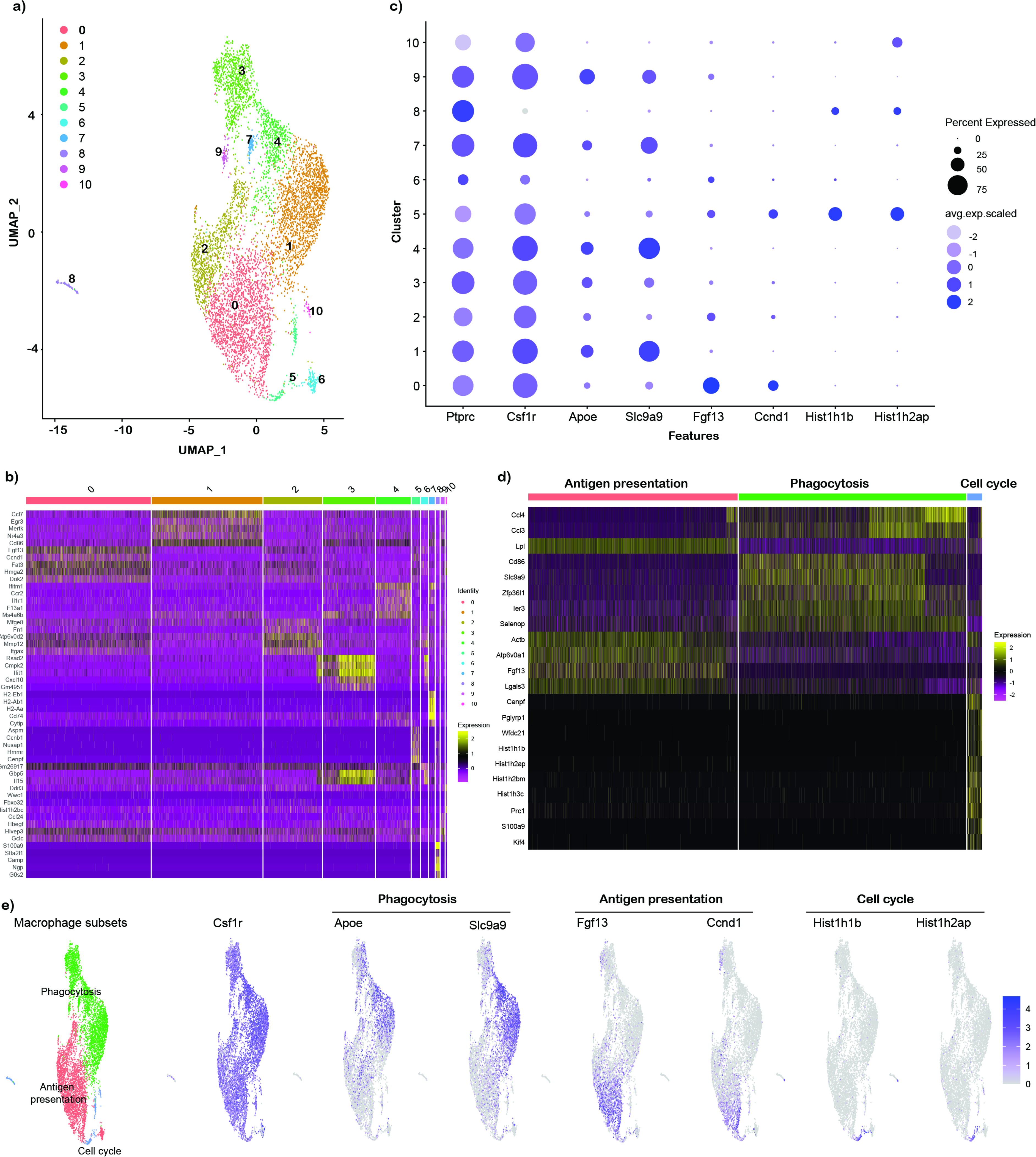
Murine bone marrow derived macrophage (BMDM) clusters identified using scRNA-Seq. FASTQ files from control BMDM (Ctrl) or exposed to *B. burgdorferi* lipoproteins (*Bb*LP) for 1 hr (LP1) or 4 hrs (LP4) were processed by CellRanger, subjected to harmony integrated UMAP. a) Total of ten clusters were identified based on the DEGs using Seurat. b) Heat map showing the top 5 genes from each cluster, showing several clusters sharing similar gene expression pattern, where the cluster information are showed on the top of the plot and the scale showing high expression (yellow) and low expression (red). c) Bubble plot showing the expression of selected genes including macrophage markers (*Ptprc*, *Csf1r*), antigen presentation markers (*Fgf13*, *Ccnd1*), phagocytosis markers (*Apoe*, *Slc9a9*) and cell cycle (*Hist1h1b*, *Hist1h2ap*). d) Heat map showing the top 7 DEGs between the clusters grouped as antigen presentation (clusters 0,2,5,6), phagocytosis (clusters 1,3,4,7,9) and cell cycle (clusters 5 and 8). e) UMAPs showing the expression pattern of select genes from each macrophage subset, with the scale at the right.

### *Bb*LP induce transcriptional reprogramming of BMDM towards a pro-inflammatory state

Harmony-integrated analysis using Seurat revealed a significant shift in the clustering of BMDMs treated with *Bb*LP at 4 hours (LP4) compared to those treated at 1 hour (LP1) and the control (Ctrl). (**Fig. 2a**). Notably, while cells in the Ctrl and LP1 groups were distributed across several clusters, most LP4 cells grouped predominantly within the phagocytosis cluster (**Fig. 2b**).

**Figure 2:**
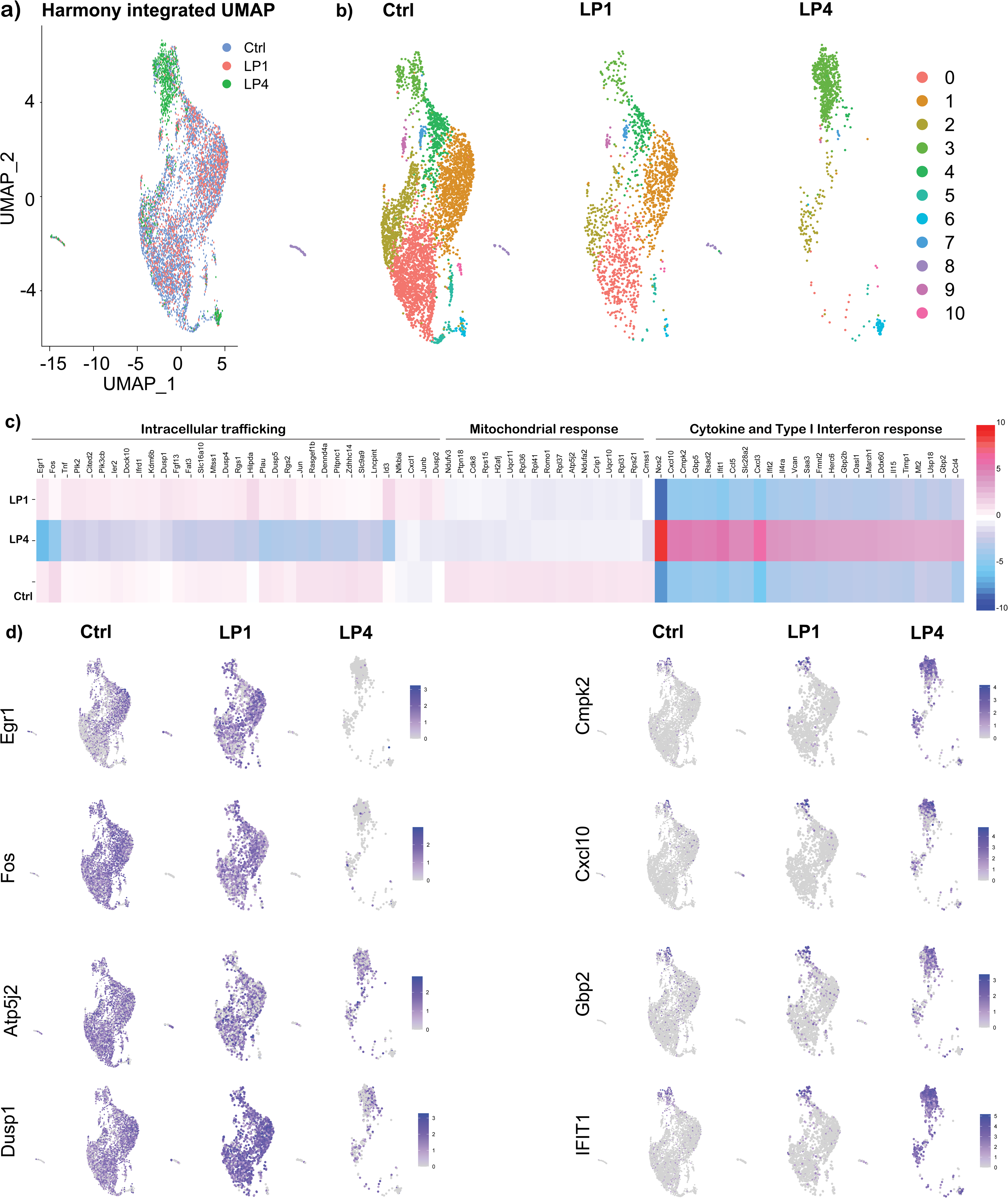
Transcriptional reprogramming of BMDMs by *Bb*LP. a) Harmony integrated UMAP displaying the cells grouped as samples, where blue dots represent cells from Ctrl, red from LP1 and green from LP4 samples. b) Individual UMAPs displaying clusters from each samples All green cells were grouped together in LP4, denoting transcriptional reprogramming in macrophages towards a pro-inflammatory phenotype. c) Heat map showing top DEGs identified from all three clusters and the functional relevance of the identified DEGs are displayed at the top of those genes as proliferation and differentiation, intracellular trafficking, mitochondrial responses, where the genes are downregulated in LP4, while cytokine and type I interferon response associated genes are upregulated in LP4, compared to Ctrl and LP1. d) UMAP plots showing differential expression of select genes between the samples.

Pseudotime analysis showed that BMDMs progress from a baseline state (control) to more activated states after 1-hour (LP1) and 4-hour (LP4) *Bb*LP treatments. Control clusters (1, 2, and 4) had early pseudotime values, indicating a mostly unstimulated state. LP1 clusters (3, 5, 6, 11, 13) showed intermediate pseudotime values, marking the initial activation response. In contrast, LP4 clusters (7, 8, 10, 12) had later pseudotime values, reflecting a more distinct response. These patterns suggest that macrophages progressively change their transcriptional profile as they respond to *Bb*LP over time (**Supplementary-Fig. 4**).

In addition, *Bb*LP treatment significantly upregulated several genes in murine BMDMs at both 1- and 4-hours post-treatment (hpt). DEG analysis revealed the upregulation of 24 genes at 1 hpi and 280 genes at 4 hpi (p<0.01) compared to unstimulated BMDMs indicating distinct transcriptional changes over time (**Supplementary-Table 2**), Comparative analysis across the three samples showed a pronounced difference in LP4, where genes involved in proliferation and differentiation (*Fos*, *Fosb*, *Rgs2*, *Egr1*, and *Egr2)*, intracellular-trafficking (*Rab7b*, *Pik2*, *Pik3cb*, and *Tnf*) and mitochondrial energy metabolism (*Atp5j2*, *Ndufv3*, *Ndufa2*, *Uqcr10*, *Uqcr11*) were downregulated. In contrast, LP4 sample exhibited upregulation of interferon response and monocyte recruitment genes (*Ifit1*, *Ifi204*, *Isg15*, *Gbp2*, *Gbp5*, *Hcar2*, *Rsad2*), cell signaling receptors (*Cd274*, *Icam1*) and inflammatory cytokine responses (*Cxcl3*, *Cxcl10*, *Ccl5*, *Il1b*, *Il15*) (**Fig. 2c**). In LP1, genes encoding several dual-specificity phosphatases such as *Dusp1*, *Dusp2*, *Dusp4*, *Dusp5* are upregulated compared to LP4 and Ctrl. In addition to these genes, receptors and signaling molecules modulated by lipoprotein ligands such as *Tlr2* and *Myd88* were also upregulated in treated BMDMs. We further used Harmony-integrated UMAP plots to analyze macrophage subset-specific expression of selected DEGs across the Ctrl, LP1, and LP4 groups. Notably, genes such as *Egr1*, *Fos*, *Id3*, and *Dusp1* were almost entirely downregulated in all clusters of LP4 compared to Ctrl and LP1. In contrast, *Cxcl10* (cytokine), *Cmpk2* (mitochondrial gene), and type-I interferon-inducible genes *Gbp2* and *Ifit1* were highly expressed in all clusters of LP4 compared to Ctrl and LP1 (**Fig. 2d**). These findings suggested that all macrophage subsets uniformly responded to *Bb*LP, likely due to the robust activation of the TLR2-MyD88 signaling cascade in BMDMs. Together, these gene expression changes reflect the transcriptional reprogramming of macrophages in response to borrelial lipoproteins, offering insights into potential targets for modulating disease outcomes. Overall, the data indicated that *Bb*LP induced significant transcriptional changes in BMDMs, skewing them toward a pro-inflammatory phenotype, as evidenced by the DEGs in LP4 and LP1 compared to Ctrl. The identified markers and the temporally constrained transcriptional changes in lipoprotein-treated macrophages provide potential therapeutic targets to modulate inflammation and pathogen clearance from reservoir hosts.

### Networks and pathways modulated by *Bb*LP in BMDMs

STRING protein network analysis revealed that most upregulated genes in LP1 and LP4 are functionally interconnected. Compared to Ctrl, mitochondrial ATP synthase genes, (*Atp5e*, *Atp5k*, *Atp5j2*); ubiquinol cytochrome C complex genes, (*Uqcrq*, *Uqcr10*, *Uqcr11)* and several ribosomal genes are downregulated in LP1 and LP4 (**Fig. 3a**), suggesting a reduction in the capacity for energy production and protein synthesis, respectively, in infected macrophages [60]. In LP1 clusters, genes encoding DUSP1/2/4/5 phosphatases are significantly upregulated compared to Ctrl and LP4 clusters (**Fig. 3b**). In LP4 clusters, several genes encoding type-1 interferon stimulated genes and signaling molecules including pro-inflammatory cytokines are significantly upregulated compared to control and LP1 clusters (**Fig. 3c**). Cellular localization analysis showed that the majority of genes upregulated in Ctrl is associated with mitochondria, LP1 is in endosomal compartments and LP4 in lysosome and sub-cellular vesicles (**Fig. 3d,e,f**). KEGG pathway analysis of the identified DEGs using InnateDB database showed that pathways such as MAPK, NOD-like receptor, Toll-like receptor, T cell receptor, RIG-I like receptor and chemokine signaling pathways are involved in LP1 and LP4 BMDMs (**Supp. Fig 5a-c**). Notably, Fc epsilon RI signaling pathway related genes are upregulated in LP1 while Fc gamma receptor mediated phagocytosis and Jak-Stat pathways are upregulated in LP4.

**Figure 3:**
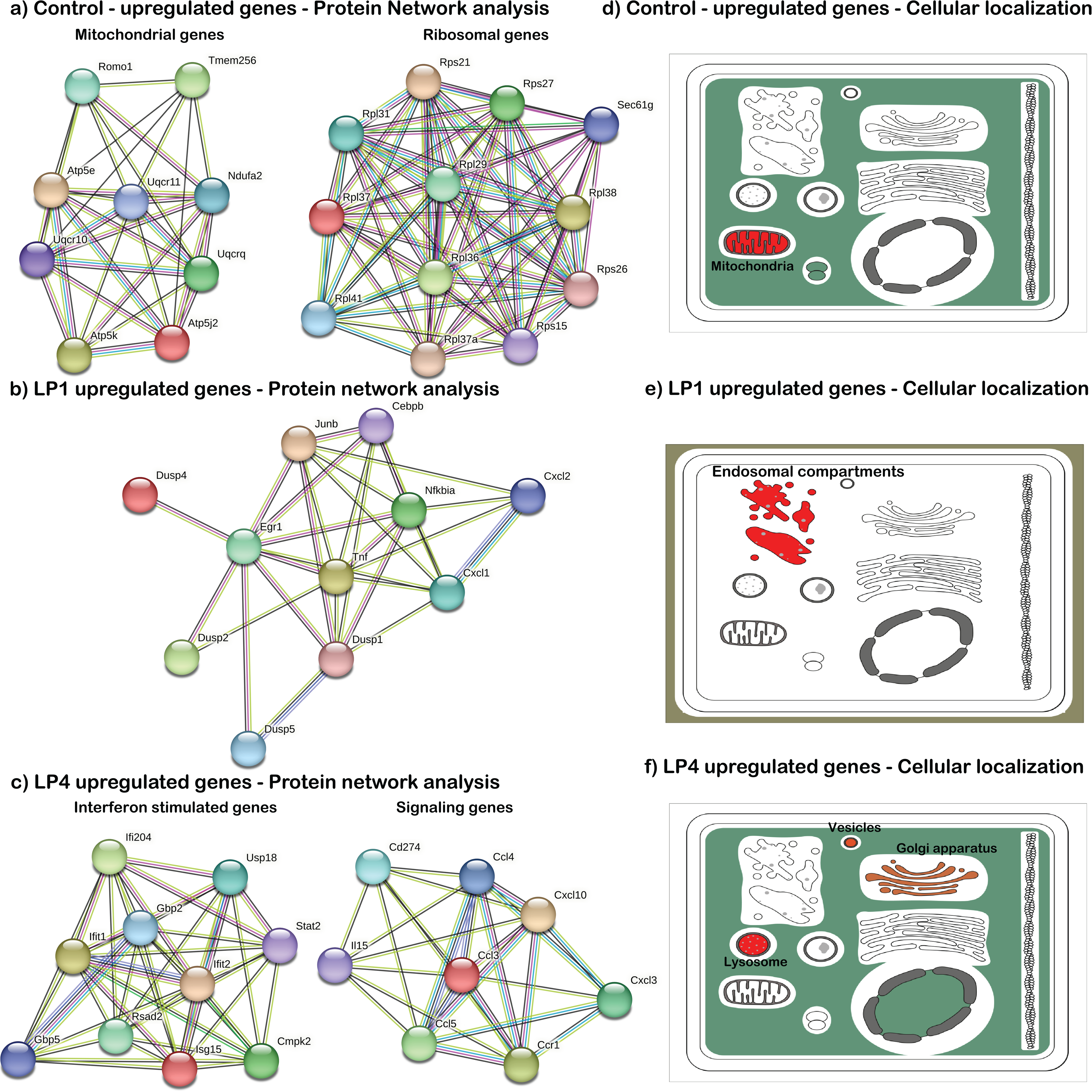
Network and cellular localization analysis of the upregulated genes in BMDMs treated with *Bb*LP. Protein network analysis of the genes that are upregulated in a) untreated BMDM (Ctrl), and BMDM treated with *Bb*LP b) at 1 hpt (LP1) and c) 4 hpt (LP4) compared to the uninfected BMDM. The lines between each circle shows the interaction between the identified genes. Proteins related to specific functions are indicated by blue rings along with the functional relevance. Graphical representation of cellular localization of the upregulated genes in d) untreated BMDM (Ctrl), and BMDM treated with *Bb*LP e) at 1 hpt (LP1) and f) 4 hpt (LP4). Red color indicate that more upregulated genes are associated with that organelle.

### Proteomics analysis

Total protein lysates of BMDMs either untreated (Ctrl) or treated with *Bb*LP (1µg/mL) for 1 (*Bb*LP1) or 4 hrs (*Bb*LP4) were subjected to unbiased mass spectrometry (MS) analysis. Two biological replicates for each sample were analyzed. Protein abundance analysis indicated that all the samples exhibited similar protein abundance (**Fig. 4a**). There were significant differential protein expression patterns between all three samples (**Fig. 4b**). 4253 individual proteins were identified from Ctrl and *Bb*LP treated samples, among which 49 shared homologies to *B. burgdorferi* proteome. After removing the low abundant non-consensus proteins, 21 proteins were identified as Borrelial proteins that are significantly expressed only in *Bb*LP treated samples but not in Ctrl BMDMs (**Fig. 4c**). The identified borrelial proteins include OspA/B/C/D, lipoproteins, Flagellar filament, and a few membrane proteins, reflecting the borrelial lipoproteins as the predominant component of borrelial subcellular fractions used in the study.

**Figure 4:**
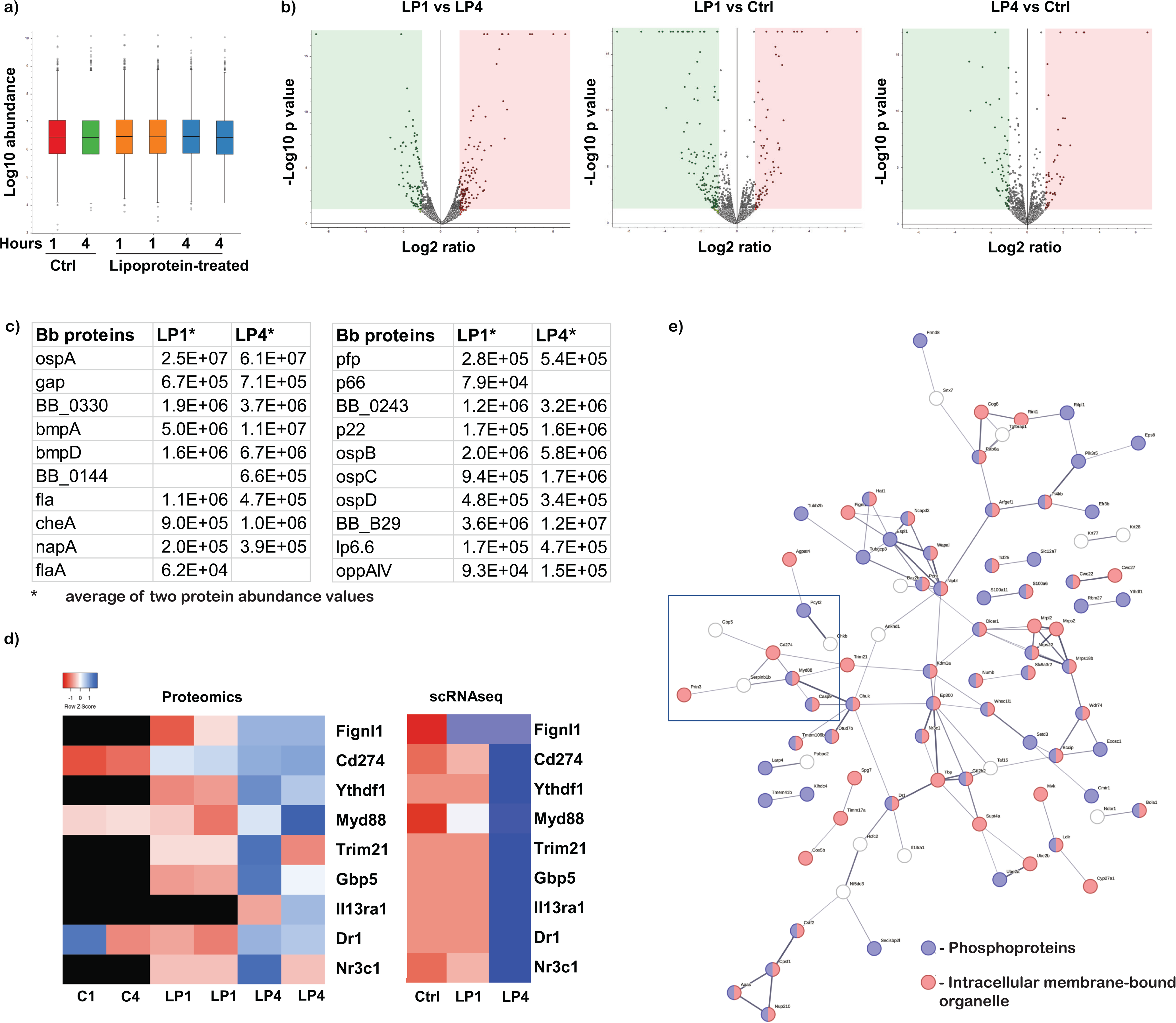
Proteomics analysis of BMDMs treated with *Bb*LPs. a) Box plot showing the normalized protein intensities of 6 protein samples extracted from BMDM, treated with lipoprotein for 1 and 4 hrs or untreated control in duplicates. B) Volcano plots visualizing the −log10 (p-value) versus the log2 fold change of proteins between the three samples with sample information mentioned at the top of each plot. Dots colored gray indicate proteins with a p-value >0.05 while red and green indicate upregulated and down regulated proteins respectively with a p-value <0.05. c) List of identified Borrelial proteins from the BMDM treated with *Bb*LP at 1 hpt (LP1) and 4 hpt (LP4) samples. d) Heat maps showing the upregulation of expression of nine consensus proteins identified from proteomics and scRNA-Seq. Scale bar represent that red indicates down regulation and blue indicates upregulation while white represent no difference in expression. Black represent absence of expression of the selected proteins. e) Protein network analysis of the upregulated proteins from the BMDM treated with *Bb*LP at 4 hpt (LP4) compared to untreated BMDM. Blue circle represents phosphoproteins and the red circles represent the proteins involved in intracellular-membrane bound organelles.

Among the 4253 total identified proteins, 4185 proteins were identified from *Mus musculus and* the abundance of each protein was compared between the six samples (2 each for untreated and 1 and 4 hpt). Low abundant proteins were removed and only proteins with abundance of over 1x10^4^ were retained for the comparative analysis. Comparative proteomics analysis between LP4 and Ctrl samples further corroborated scRNA-Seq findings such as activation of proteins involved in the MyD88/CD274(PD-L1) axis as well as select proteins GBP5, IL13RA1, FIGNL1, NR3C1, among others (**Fig. 4d**). The glucocorticoid receptor (GR, encoded by *Nr3c1*), which translocates to the nucleus upon binding to a ligand to regulate gene transcription, regulates several genes including PD-L1, while GBP5, IL13RA1, FIGNL1 are regulated by interferon [61–64]. Although the expression of fidgetin-like-1 (FIGNL1) does not directly interact with the immune system, its role in cell division and motility could influence macrophage recruitment to the site of infection. Consistent with the scRNA-Seq analysis, several kinases and phosphatases were upregulated in LP4 samples. Protein network analysis showed that majority of the identified proteins are phosphoproteins localized to the intracellular membrane bound organelles (**Fig. 4e**), suggesting a role for these phosphoproteins following treatment of macrophages with LP4. Moreover, several upregulated proteins in LP4 were consistent with the scRNA-Seq cellular localization analysis, where most of the upregulated genes are localized in the sub-cellular vesicles and lysosome (**Fig 3f**).

### *Bb*LP transiently upregulates DUSP1 in BMDMs

Quantitative RT-PCR analysis of BMDMs treated with 1 and 0.1 µg of *Bb*LP showed increased transcriptional levels of *Dusp1* at 1 hr and the expression levels were subsequently reduced at 4 and 24 hrs (**Fig. 5a**). Immunoblot analysis also showed increased levels of DUSP1 at 1 and 2 hpt with a marked reduction at 48 hrs correlating with the transcriptional data (**Fig. 5b**). To delineate the role of IκB kinase (IKK) in controlling DUSP1 protein expression and determine the regulatory role of DUSP1 during *Bb*LP treatment, BMDMs were treated with Bay 11-7082 (Ikb kinase inhibitor) or BCI (DUSP 1/6 inhibitor), respectively for 1 hr prior to treatment with *Bb*LP. Both IKK and DUSP1 inhibitors significantly reduced the DUSP1 levels in *Bb*LP-treated BMDM, compared to *Bb*LP-treated and *E. coli* LPS treated BMDMs for 1 hr (**Fig. 5c**). Furthermore, confocal microscopy showed that DUSP1 levels were higher in *Bb*LP-treated BMDMs compared to the controls (**Fig. 5d**), while BCI treated BMDMs had reduced DUSP1 expression irrespective of *Bb*LP treatment, indicating that BCI treatment reduced the DUSP1 protein levels in BMDM. Taken together, these results showed that DUSP1 expression was induced during early stages of treatment of BMDMs with *Bb*LP and that its levels could be reduced using a specific inhibitor, BCI.

**Figure 5:**
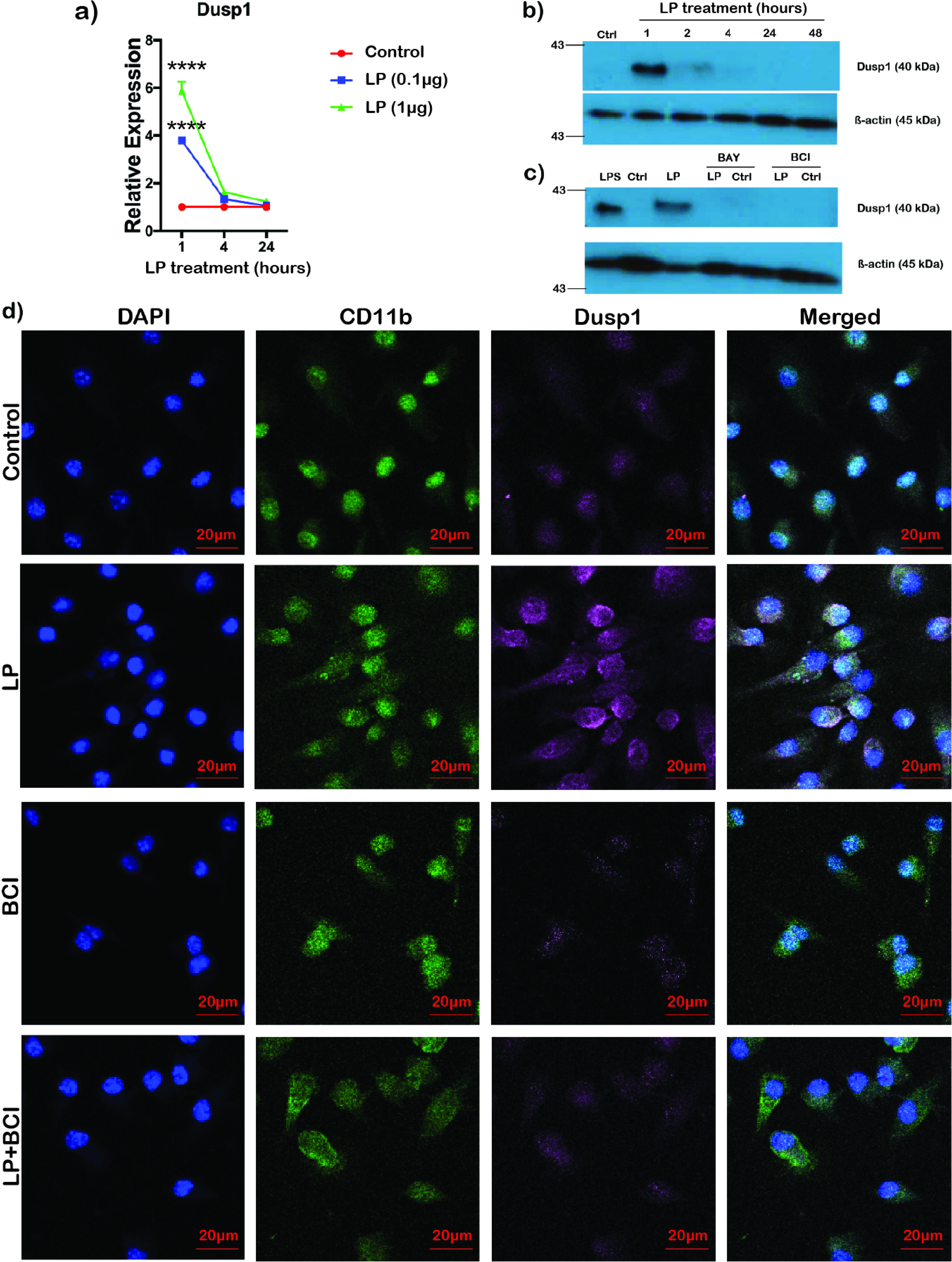
Transient upregulation of Dusp1 in BMDMs treated with *Bb*LP. a) Line graph showing the kinetics of relative expression of *Dusp1* gene in BMDM treated with *Bb*LP (1 and 0.1 µg) at 1, 4 and 24 hrs determined by qPCR analysis. Dusp1 protein expression detected by western blotting b) from LP (1 µg) treated BMDM at 1, 2, 4, 24 and 48 hpi compared to the uninfected BMDM. c) from BAY and BCI pre-treated BMDM, followed by *Bb*LP (1 µg) treated for 1 hpi. *E. coli* LPS was used as positive control and uninfected BMDM was used as negative control. Mouse actin was used as the normalization control for both the blots. d) Confocal microscopy of the colocalization of DUSP1 (Purple) and CD11b (Green) in BMDM at 1 hpi with and without LP treatment (Control). BMDM were either pre-treated with BCI inhibitor or untreated for 1 hr before LP treatment. Data are presented as mean ± standard error of the mean (SEM). Statistical significance between samples is indicated as follows: *p < 0.05, **p < 0.01, ***p < 0.001, ****p < 0.0001.

### NLRP3-inflammasome pathway associated genes are upregulated in BMDMs by *Bb*LP

The scRNA-Seq analysis revealed that NLRP3-inflammasome pathway-associated genes, including *Nlrp3*, *Cxcl1*, *Cxcl2*, *Ccrl2*, and *Il1f9*, were significantly upregulated in BMDMs at 1 and 4 hpt with *Bb*LP. To confirm this upregulation, BMDMs were treated with 1 and 0.1 µg/mL of *Bb*LP for 1, 4, and 24 hours. Transcriptional levels of *Nlrp3*, *Cxcl1*, *Cxcl2*, *Ccrl2*, and *Il1f9* were significantly upregulated in BbLP-treated BMDMs, irrespective of dose, compared to untreated BMDMs at all timepoints, with *Actb* serving as a control (**Fig. 6a; supplementary Table 3**).

**Figure 6:**
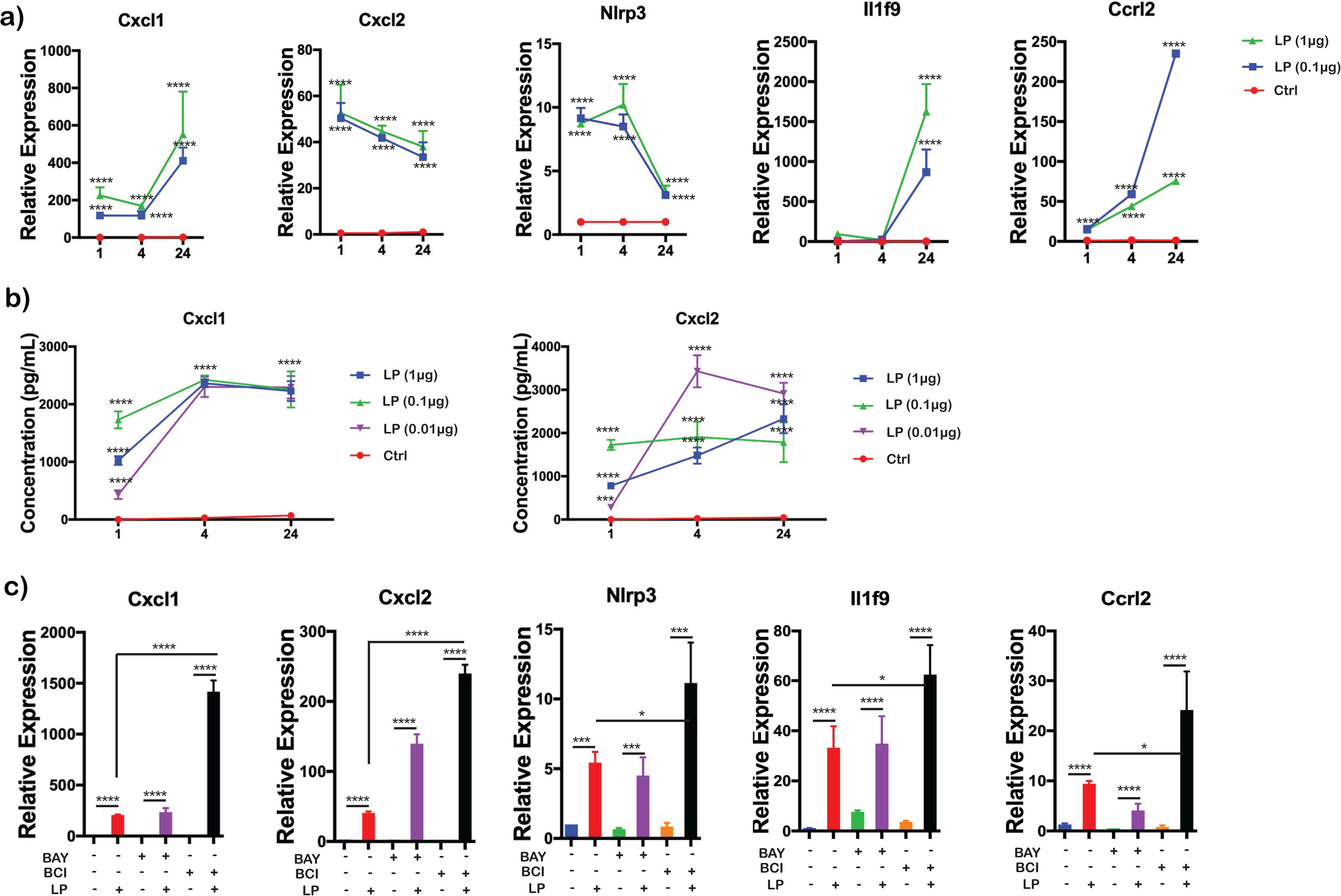
Modulation of Nlrp3-associated genes by IKK and DUSP1/6 inhibitors in BMDMs treated with *Bb*LP. a) Line graphs showing the kinetics of relative expression of *Cxcl1*, *Cxcl2*, *Nlrp3*, *Il1f9* and *Ccrl2* genes in BMDM during *Bb*LP treatment (1 and 0.1 µg) at 1, 4 and 24 hrs determined by RT-PCR analysis. b) Line graphs showing the levels of cytokines Cxcl1 and Cxcl2 from the supernatant of LP-treated (1, 0.1 and 0.01 µg) and untreated BMDM quantified by ELISA. c) Bar graphs showing the relative expression of *Cxcl1*, *Cxcl2*, *Nlrp3*, *Il1f9* and *Ccrl2* genes in BMDM during *Bb*LP treatment (1 µg) pre-treated with or without BAY and BCI inhibitors, determined by RT-PCR analysis. Data are presented as mean ± standard error of the mean (SEM). Statistical significance between samples is indicated as follows: *p < 0.05, **p < 0.01, ***p < 0.001, ****p < 0.0001.

Additionally, BMDMs were treated with 1, 0.1, 0.01, and 0 µg/mL of *Bb*LP for 1, 4, and 24 hours and ELISA was performed to measure CXCL1 and CXCL2 levels in the culture supernatants. Both CXCL1 and CXCL2 cytokine levels were significantly elevated across all *Bb*LP doses, with no expression detected in untreated BMDMs (**Fig. 6b**). Notably, CXCL1 and CXCL2 expression was significantly higher at 4 and 24 hours compared to 1 hour, indicating sustained *Bb*LP-induced cytokine production in BMDMs.

### IKK and DUSP1/6 inhibitors modulate *Bb*LP induced expression of NLRP3-associated genes

To understand the roles of IKK and dual specificity phosphatases in regulating the expression of NLRP3-associated genes in response to *Bb*LP treatment, BMDMs were treated with BAY or BCI inhibitors prior to treatment with *Bb*LP (0.1 µg/mL). BMDMs treated with *Bb*LP with no inhibitors were maintained as controls. At 4 hpt, RNA was isolated and RT-PCR analysis was performed. The expression of *Nlrp3*, *Cxcl1*, *Cxcl2*, *Ccrl2*, and *Il1f9* genes was significantly upregulated in *Bb*LP-treated BMDMs in the presence of BCI inhibitor, compared to *Bb*LP-treated BMDMs without inhibitors, suggesting that DUSP1 is necessary to control the expression of NLRP3-pathway associated genes during *Bb*LP treatment (**Fig. 6c**). No significant changes were observed in the expression of these genes suggesting that inhibition of IKK by BAY does not affect lipoprotein-stimulated expression of NLRP3-associated genes (**Fig. 6c**). Together, these results indicate that dual specificity phosphatases negatively regulate the *Bb*LP-induced expression of NLRP3-associated genes in BMDM.

### *Bb*LP induce time-dependent changes in inflammatory cytokines

scRNA-Seq analysis indicated that *Bb*LP upregulated several inflammatory cytokine genes, including *Ccl5* and *Il1b*, in the LP4 sample (**Fig. 2**). To validate scRNA-Seq data, the upregulation of these genes was confirmed by RT-PCR using BMDMs treated with *Bb*LP (0.1 and 1 µg/mL). As shown in Fig. 7a, the expression levels of *Ccl5* and *Il1b* were significantly upregulated in a time-dependent but dose-independent manner, compared to BMDM not treated with *Bb*LP (**Fig. 7a**). In addition, inhibitors of IKK and DUSP significantly reduced the levels of *Ccl5* and *Il1b* transcriptionally compared to BMDMs treated with *Bb*LP alone (**Fig. 7b**). Moreover, BCI was relatively less effective in reducing the expression of CCL5 in *Bb*LP-treated BMDMs compared to the IKK inhibitor BAY. These results suggest that IKK and DUSP1 positively regulate the expression of *Ccl5* and *Il1b* in BMDM following *Bb*LP treatment.

**Figure 7:**
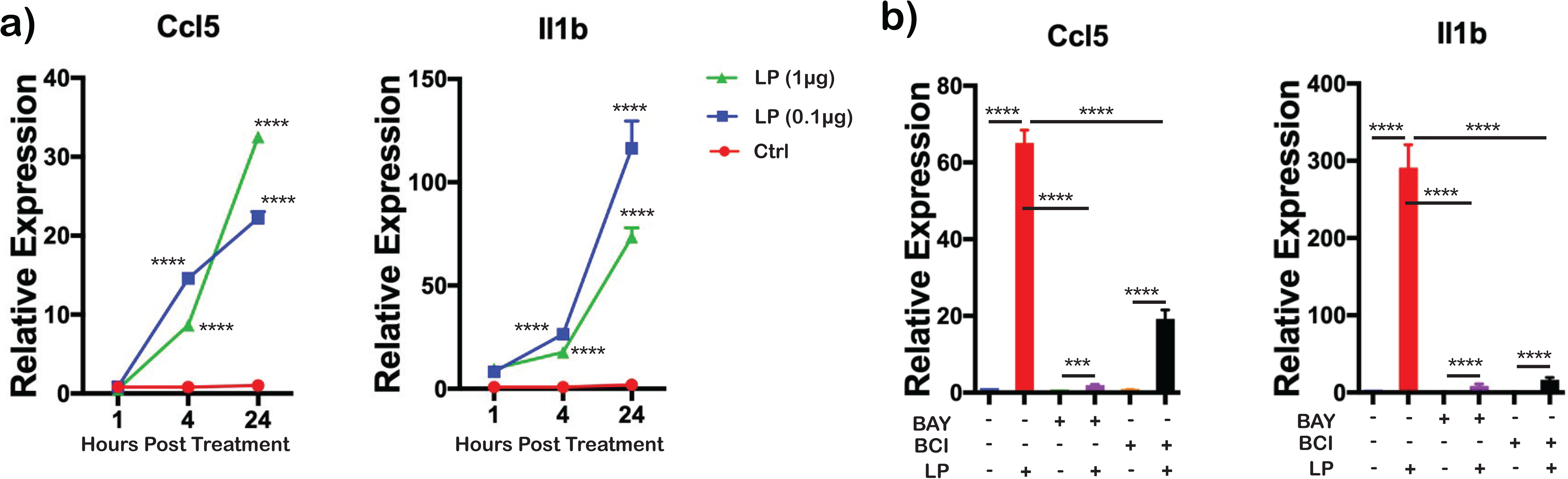
Modulation of pro-inflammatory cytokines by *Ccl5* and *Il1b* in BMDMs treated with *Bb*LP. a) Line graphs showing the time-dependent increase in the expression of *Ccl5* and *Il1b* in BMDM during *Bb*LP treatment (1 and 0.1 µg) at 1, 4 and 24 hrs compared to the uninfected BMDM determined by RT-PCR analysis. b) Bar graphs showing the relative expression of *Ccl5* and *Il1b* genes in BMDM during *Bb*LP treatment (1 µg) pre-treated with or without BAY and BCI inhibitors, determined by RT-PCR analysis. Data are presented as mean ± standard error of the mean (SEM). Statistical significance between samples is indicated as follows: *p < 0.05, **p < 0.01, ***p < 0.001, ****p < 0.0001.

### *Bb*LP and Pam3CSK4 dependent modulation of select cytokines

We further dissected the ability of the structural features of borrelial lipoproteins in modulating the levels of select cytokines via IKK/DUSP1 dependent manner in comparison to TLR2 ligand Pam3CSK4 which is a mimic of the post-translational modification of borrelial lipoproteins [65]. As shown in Fig. 8a, levels of cytokines CXCL1, CXCL2, CCL5 and IL1B were elevated in culture supernatants of BMDMs treated with *Bb*LP or Pam3CSK4 at 24 hpt indicating that these ligands activate a comparable initial cytokine response through TLR2 signaling (**Fig. 8a,b**). BMDMs treated with IKK inhibitor BAY exhibited no significant changes in CXCL1, CXCL2, CCL5 and IL1B levels between unstimulated and *Bb*LP stimulated cells suggesting that the effects of *Bb*LP at the concentrations used were unable to overcome IKK inhibition. However, treatment of Pam3CSK4 was able to overcome IKK inhibition by BAY resulting in significantly higher levels of CXCL1, CXCL2 and IL1B while there was no such change in the levels of CCL5. In the presence of DUSP1 inhibitor BCI, CXCL1, CXCL2 and CCL5 were significantly elevated in the presence of *Bb*LP with no such significant change in the levels of Il1b. However, the addition of Pam3CSK4 was able to overcome the effects of BCI resulting in significantly higher levels of CXCL1, CXCL2, CCL5 and IL1B suggesting the possibility of DUSP1 independent mechanisms of stimulation of these cytokines (**Fig. 8a, b**). Alternatively, it is also possible that concentration/structural features of Pam3CSK4 as a purified PAMP was more effective than as part of purified *Bb*LP in interacting with TLRs.

**Figure 8:**
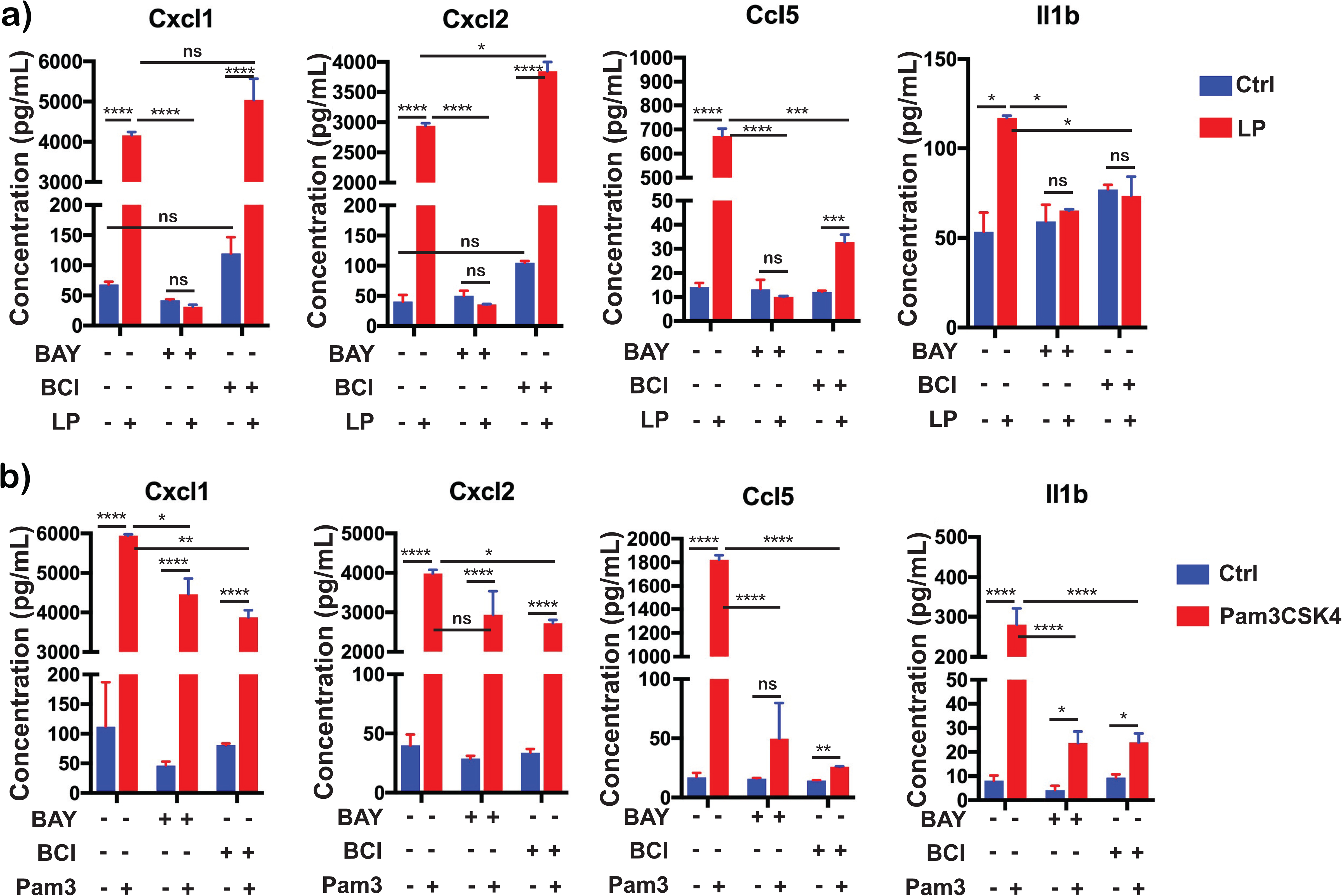
Differential cytokines expression induced in BMDMs by *Bb*LP and Pam3CSK4 pre-treated with IKK and Dusp1/6 inhibitors. Bar graphs showing the levels of cytokines, CXCL1, CXCL2, CCL5 and IL1B determined from the supernatant of BMDM pre-treated with or without inhibitors of IKK (BAY) and Dusp1/6 (BCI) followed by the treatment of a) LP (1 µg) and b) Pam3CSK4 (1µg) at 24 hpi. Data are presented as mean ± standard error of the mean (SEM). Statistical significance between samples is indicated as follows: *p < 0.05, **p < 0.01, ***p < 0.001, ****p < 0.0001.

### Type I interferon stimulated genes are upregulated by *Bb*LP in BMDM

scRNA-Seq analysis indicated that *Bb*LP upregulated type I interferon stimulated genes, commonly referred to as ISGs (Interferon-Stimulated Genes) in BMDM at 4 hpt but not at 1 hpt (Fig 2). To further validate these findings, BMDMs were treated with *Bb*LP (0.1 and 1 µg/mL) and RNA was isolated from BMDMs at 1, 4 and 24 hpi. RT-PCR analysis showed upregulation of transcriptional levels of select ISGs such as *Gbp2*, *Gbp5*, *Ifit1*, *Isg15*, *Hcar2* and *Rsad2* in a time-dependent manner with both concentrations of *Bb*LP compared to untreated controls (**Fig. 9a**).

**Figure 9:**
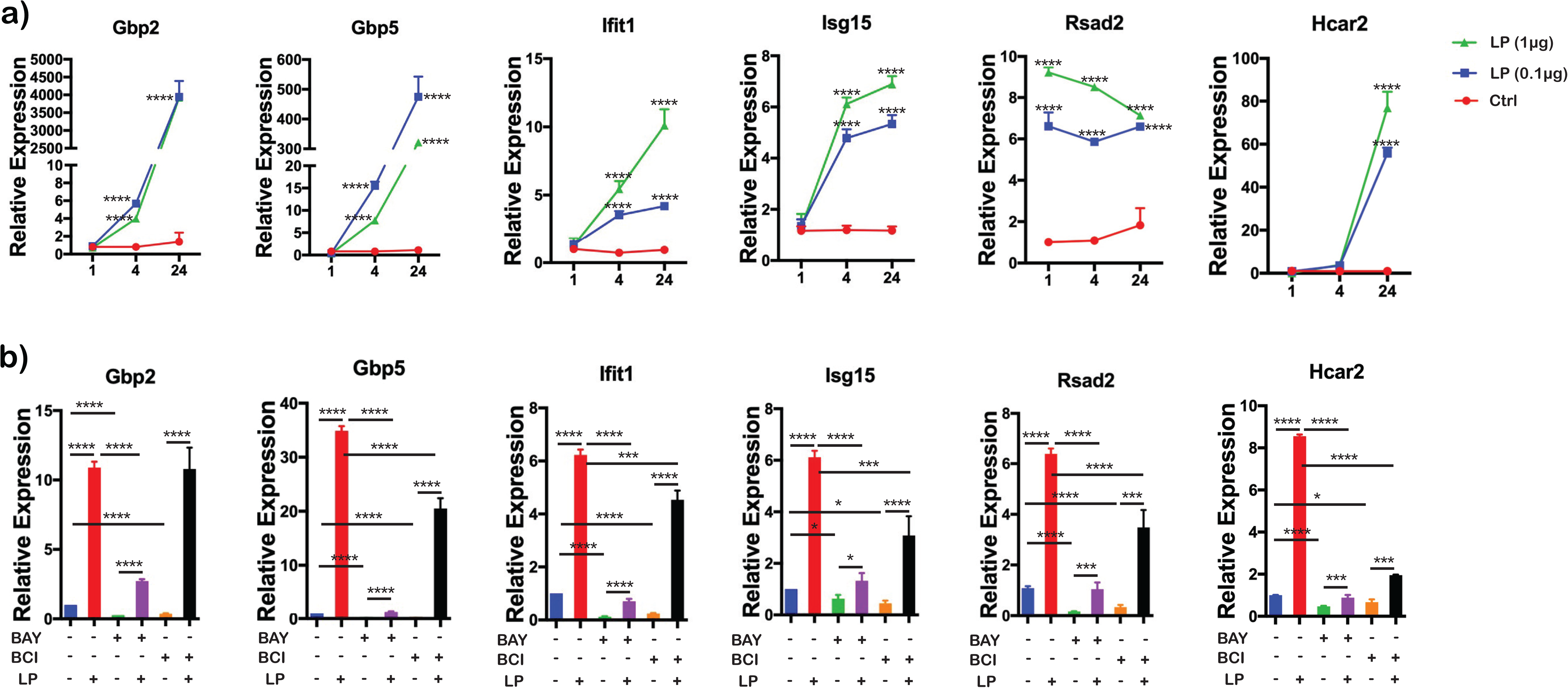
Modulation of IRF pathway genes in BMDMs by *Bb*LP. a) Line graphs showing the kinetics of relative expression of *Gbp2*, *Gbp5*, *Ifit1*, *Isg15*, *Rsad2* and *Hcar2* genes in BMDM during *Bb*LP treatment (1 and 0.1 µg) at 1, 4 and 24 hrs determined by RT-PCR analysis. b) Bar graphs showing the relative expression of *Gbp2*, *Gbp5*, *Ifit1*, *Isg15*, *Rsad2* and *Hcar2* genes in BMDM during *Bb*LP treatment (1 µg) pre-treated with or without BAY and BCI inhibitors, determined by RT-PCR analysis. Data are presented as mean ± standard error of the mean (SEM). Statistical significance between samples is indicated as follows: *p < 0.05, **p < 0.01, ***p < 0.001, ****p < 0.0001.

While IKK inhibitor BAY significantly inhibited ISGs tested, compared to *Bb*LP alone, this inhibition was not saturating as there was some induction in BAY plus *Bb*LP treated BMDMs compared to BAY treated cells. This was also true for DUSP inhibitor BCI although the levels of the tested ISGs were significantly higher in the presence of *Bb*LP (**Fig. 9b**). These observations suggest that the IKK dependent effects on ISGs are pronounced compared to DUSP1 dependent effects.

### *Bb*LP upregulates PD-L1 expression in BMDMs

The scRNA-Seq and proteomics data sets revealed that *Cd274* gene encoding PD-L1 was upregulated in *Bb*LP infected BMDMs (**Fig. 4d**). To further validate these findings, BMDMs treated with *Bb*LP (0.1 and 1 µg/mL), stained using anti-PD-L1 antibody at 1 and 24 hpt, followed by flow cytometry analysis. Compared to control BMDMs, *Bb*LP significantly increased the surface expression of PD-L1 (PD-L1^hi^ population) at 24 hpt (**Fig. 10a-c**). No significant differences were observed between *Bb*LP treated and control BMDMs at 1 hpt. Further, to understand the involvement of DUSP1 in regulating the PD-L1 expression in response to *Bb*LP, BMDMs were first treated with BCI inhibitors followed by exposure to *Bb*LP. At 24 hpt, flow cytometric analysis revealed that there was a significant reduction in PD-L1 expression in the BCI-treated BMDMs with or without *Bb*LP treatment (**Fig. 10d-f**) indicating that *Bb*LP induced expression of PD-L1 in macrophages is regulated by DUSP1.

**Figure 10:**
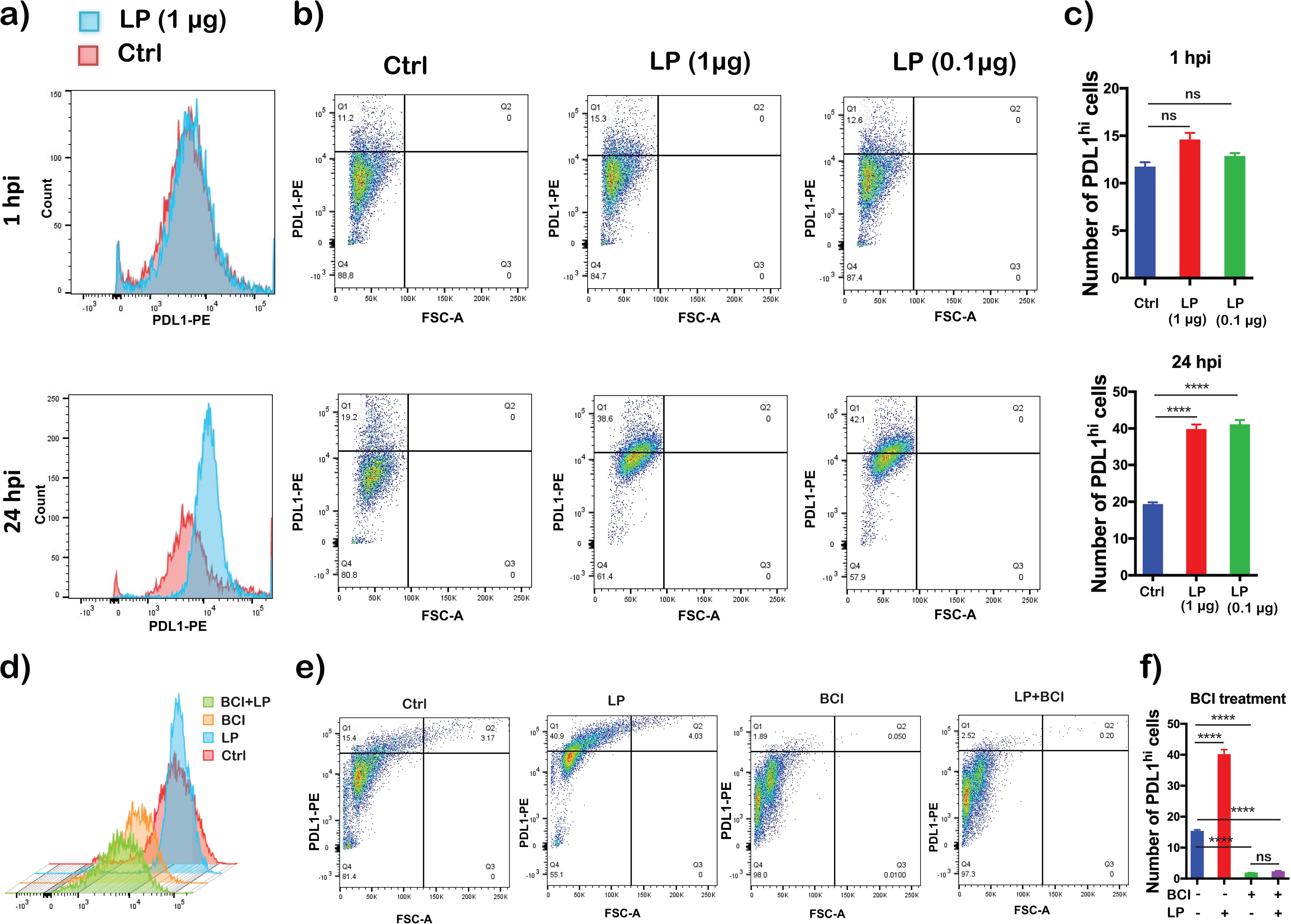
Flow cytometry analysis depicting expression levels of PD-L1 in *Bb*LP treated BMDMs. Expression pattern of PD-L1 in BMDM with or without LP treatment a) Histogram, b) Dot-plot, c) Bar-graph showing the percentage of PD-L1^hi^ cells. Expression pattern of PD-L1 in BCI pre-treated or untreated BMDM before LP treatment. d) Histogram, e) Dot-plot, f) Bar-graph showing the percentage of PD-L1^hi^ cells. Data are presented as mean ± standard error of the mean (SEM). Statistical significance between samples is indicated as follows: *p < 0.05, **p < 0.01, ***p < 0.001, ****p < 0.0001.

Total cell lysates of BMDMs treated with *Bb*LP for 1, 2, 4, 24 and 48 hrs were analyzed for PD-L1 levels by immunoblot analysis using commercially available anti-PD-L1 antibodies. PD-L1 levels were higher at 24 and 48 hpt compared to earlier time points (**Fig. 11a**), correlating with flow-cytometric analysis. Moreover, Pam3CSK4 treated cells induced PD-L1 activation similar to *Bb*LP at 24 hpi in BMDMs. These observations validated the multi-omics finding shown in Fig. 4d, that *Bb*LP induced significantly higher PD-L1 expression in BMDMs. Confocal image analysis also indicated elevated intensity of PD-L1 on LP treated BMDMs co-localizing with the CD11b, while the PD-L1 levels were reduced in BCI-treated BMDMs compared to *Bb*LP treated BMDMs (**Fig. 11b**). Together, these observations demonstrated that PD-L1 surface expression is increased in response to borrelial lipoproteins and that DUSP1 inhibitor, BCI reduces the *Bb*LP induced PD-L1 expression on the BMDM.

**Figure 11:**
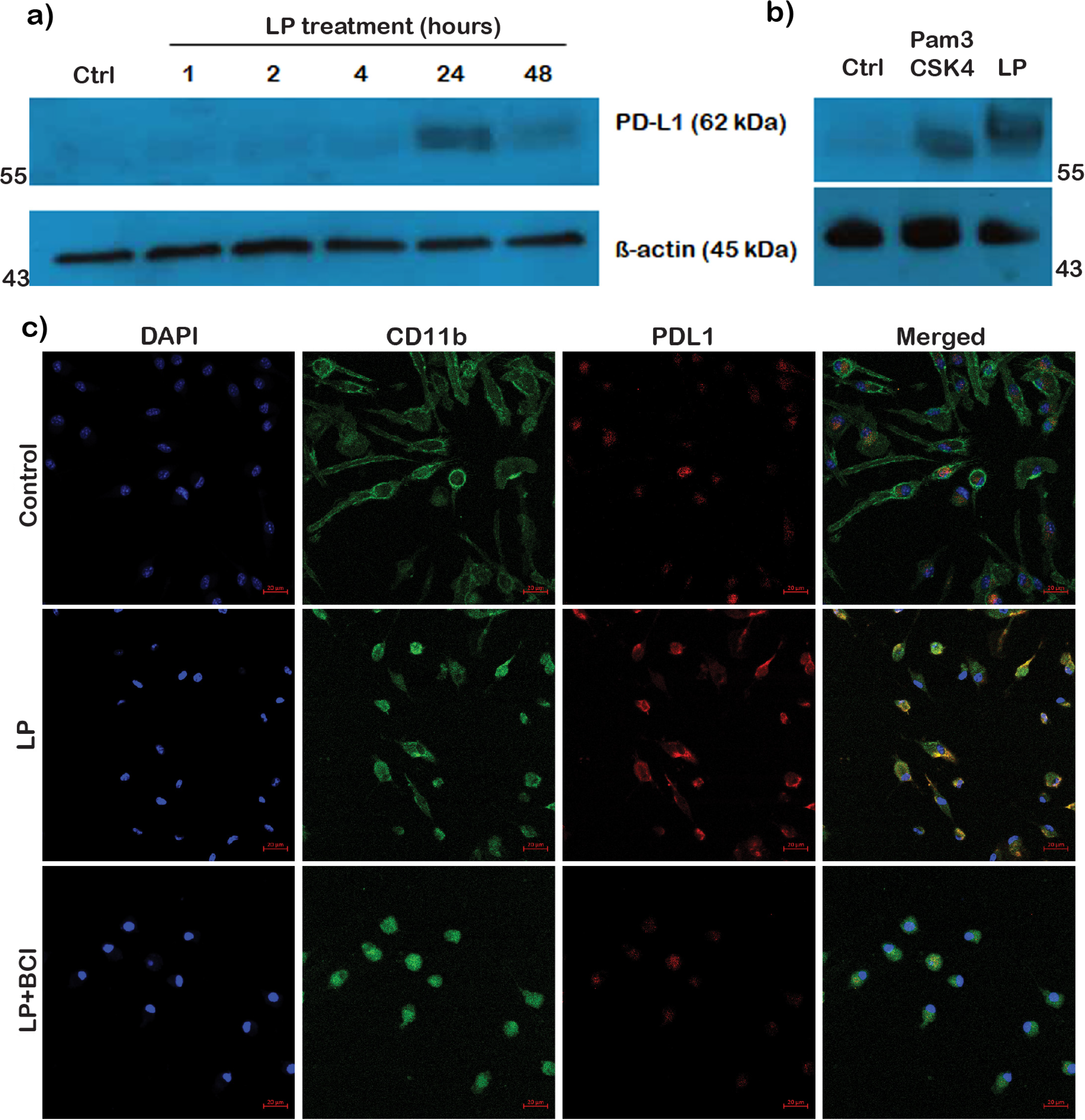
PD-L1 expression kinetics in BMDMs with *Bb*LP treatment. Western blot analysis showing the expression of PD-L1 in a) BMDM treated with *Bb*LP at 1, 2, 4, 24 and 48 hpt and b) BMDM treated with Pam3CSK4 and *Bb*LP at 24 hpt. Untreated BMDM was maintained as control and ß-actin was maintained as normalization control. c) Co-localization of PD-L1 (Red) surface expression with CD11b (Green) and nuclear stain DAPI (Blue) in BMDM after 24 hrs with or without *Bb*LP treatment and pre-treated with BCI.

### MyD88 is necessary for *Bb*LP induced NF-κB & IRF responses in macrophages

*Bb*LP upregulated NF-κB-regulated genes (**Fig. 6**) and ISGs through IRF-regulated expression in murine BMDMs by activating IKK (**Fig. 9**). To specifically show NF-κB and ISG activation by *Bb*LP in macrophages, we used the THP1 dual-reporter human monocyte cell line. This cell line includes a luciferase reporter for ISG activation and a secreted embryonic alkaline phosphatase (SEAP) reporter for NF-κB activity, allowing simultaneous measurement of both pathways by monitoring luciferase and SEAP activity, respectively [66]. THP1 cells, differentiated into macrophages using PMA, were treated with *Bb*LP, while cells treated with Pam3CSK4 served as a positive control and bovine serum albumin (BSA) as a negative control. At 24 hpi, SEAP was measured from the culture supernatants to assess NF-κB activity. THP1 cells treated with *Bb*LP showed significant SEAP activity at all tested concentrations, indicating that *Bb*LP induces NF-κB activation in THP1-derived macrophages, even at doses as low as 1 ng/mL (**Fig. 12A**). Similarly, supernatants from cells treated with 1 ng/mL of TLR2 agonist, Pam3CSK4, also had significant SEAP activity. These results indicated that *Bb*LP induces TLR2 mediated NF-κB activity in human macrophages derived from THP1 monocyte like cell line consistent with what has been observed with the murine macrophages.

**Figure 12:**
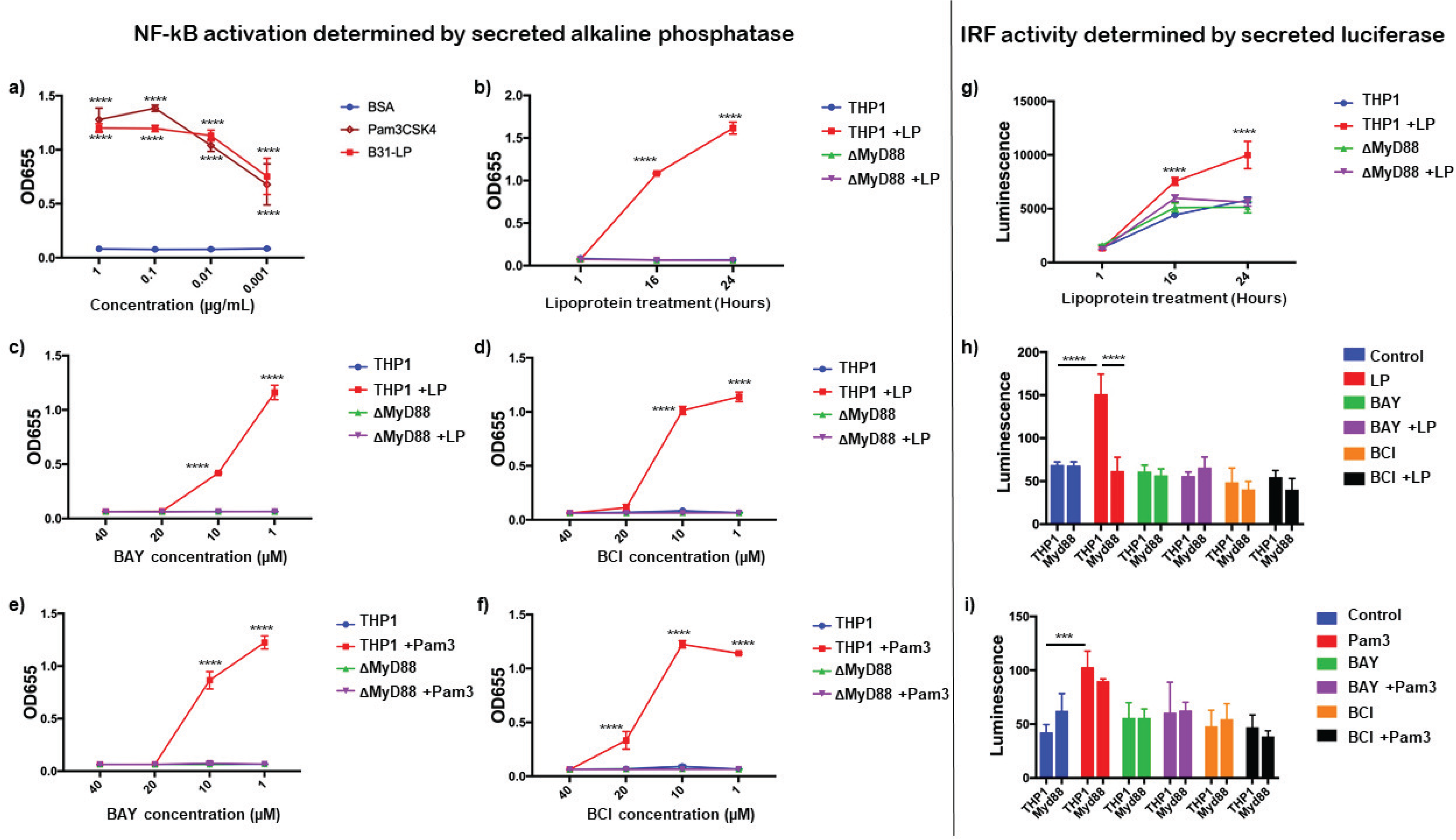
Activation of NFkB and IRF pathways in human THP1 dual reporter cells by *Bb*LP. NF-kB activation was determined by the secreted alkaline phosphatase: a) SEAP activity in THP1 cells treated with different concentrations of BSA, Pam3CSK4 and LP at 24 hrs. Kinetics of LP-induced SEAP activity in THP1 and MyD88 KO reporter cells, b) at 1, 16 and 24 hrs, and cells pre-treated with different concentrations of c) BAY inhibitor and d) BCI inhibitor. Kinetics of Pam3CSK4 induced SEAP activity in THP1 and MyD88 reporter cells pre-treated with different concentrations of e) BAY inhibitor and f) BCI inhibitor. IRF activity was determined by the secreted luciferase: g) kinetics of *Bb*LP-induced luminescence at 1, 16 and 24 hours post treatment. Bar graphs showing the luminescence induced by h) *Bb*LP and i) Pam3CSK4 in THP1 and MyD88 KO THP1 reporter cells pre-treated with BAY or BCI inhibitors, while untreated cells were maintained as control. Data are presented as mean ± standard error of the mean (SEM). Statistical significance between samples is indicated as follows: *p < 0.05, **p < 0.01, ***p < 0.001, ****p < 0.0001.

scRNA-Seq analysis and proteomics data sets revealed the upregulation of MyD88 in macrophages following *Bb*LP treatment (**Fig. 4d**). To understand the role of MyD88 in inducing NF-κB activity in macrophages, THP1 and MyD88 KO dual reporter cell lines were treated with 0.1 µg/mL *Bb*LP and SEAP activity was significantly high at 16 and 24 hrs compared to the untreated THP1 cells. However, in MyD88 KO-THP1 cells, no significant SEAP activity was observed at all the timepoints tested irrespective of the concentration of *Bb*LP (**Fig 12b**). These findings showed that *Bb*LP induced NF-κB pathway in THP1 cells via MyD88-dependent pathway consistent with previous findings [39, 45, 67].

In order to connect the effects of *Bb*LP on ISGs in human macrophages, we determined luciferase activity that tracks the IRF pathway using THP1 dual reporter cell lines [66]. The luciferase activity was significantly higher in the *Bb*LP treated THP1 cells at 16 and 24 hpi, compared to the untreated THP1 cells (**Fig 12c**). However, there was no significant change in the luciferase activity in the *Bb*LP treated MyD88 KO-THP1 cells, compared to untreated MyD88 KO cells. These results indicated that *Bb*LP treatment induces ISG54-regulated ISGs in human macrophages, via MyD88-dependent pathway.

### DUSP1 differentially regulate NF-κB pathway in *Bb*LP and Pam3CSK4 treated THP1 macrophages

To understand the involvement of IKK and DUSP1 in regulating NF-κB and IRF pathways in human macrophages induced by borrelial lipoproteins, THP1-dual reporter cells were treated with different concentrations (40, 20, 10 and 1 µM) of BAY and BCI inhibitors for 1 hr prior to treatment with *Bb*LP (0.1 µg/mL). Secreted SEAP and Luciferase activities were determined as absorbance (OD_655_) and luminescence respectively, from culture supernatants at 24hr. Both BAY and BCI inhibitors completely inhibited *Bb*LP-induced SEAP activity at 20 and 40µM (**Fig. 12d, e**). *Bb*LP-induced Luciferase activity was also inhibited by BAY and BCI inhibitors at 20 µM in THP1 cells (**Fig. 12f**). Together, the results suggested that both IKK kinases and DUSP1 phosphatase regulate the NF-κB and ISG response in macrophages following treatment with *Bb*LP.

In order to determine if the inhibition of NF-κB/ISG activity by BAY and BCI inhibitors in macrophages is Tlr2-specific, we stimulated THP1 and MyD88-KO cells with Pam3CSK4 (10 ng/mL). Similar to *Bb*LP, levels of SEAP and Luciferase activity induced by Pam3CSK4 in THP1 cells were significantly reduced in the presence of 20µM of BAY (**Fig. 12g,i**) and BCI inhibitors (**Fig. 12h,i**). As expected, MyD88 KO did not show any significant difference following treatment with inhibitors or Pam3CSK4 stimulation, compared to untreated THP1 (**Fig. 12g-i**). These results demonstrated that IKK kinase and DUSP1/6 inhibitors significantly blocked the activation of NF-κB pathway during *Bb*LP and Pam3CSK4 stimulation in macrophages. This indicated that *Bb*LP induced NF-κB/IRF pathway in macrophages through MyD88-dependent TLR2 activation and are primarily regulated by IKK kinases and DUSP1 phosphatase.

### *Bb*LP modulates mitochondrial oxidative stress response pathways in THP1 macrophages

*Bb*LP treatment downregulated several mitochondrial genes in BMDM (Fig 3a), while mitochondrial antioxidant gene *Sod2* and a mitochondria-associated kinase cytidine monophosphate kinase 2, *Cmpk2* were upregulated, suggesting *Bb*LP modulate mitochondrial oxidative stress pathways. To validate scRNA-Seq findings on the effects of *Bb*LP on the transcriptional levels of *Cmpk2* and *Sod2* (**Fig. 3c**), THP1 cells were treated with *Bb*LP for 1, 2, 4, 16, 24, and 48 hours. Expression levels of *Cmpk2* gene were significantly increased in *Bb*LP treated THP1 cells at 16 hpi and gradually decreased at 24 and 48 hpt (**Fig. 13a**), whereas *Sod2* expression was significantly increased over time in treated macrophages from 4 to 48 hpt compared to control macrophages (**Fig. 13b**).

**Figure 13:**
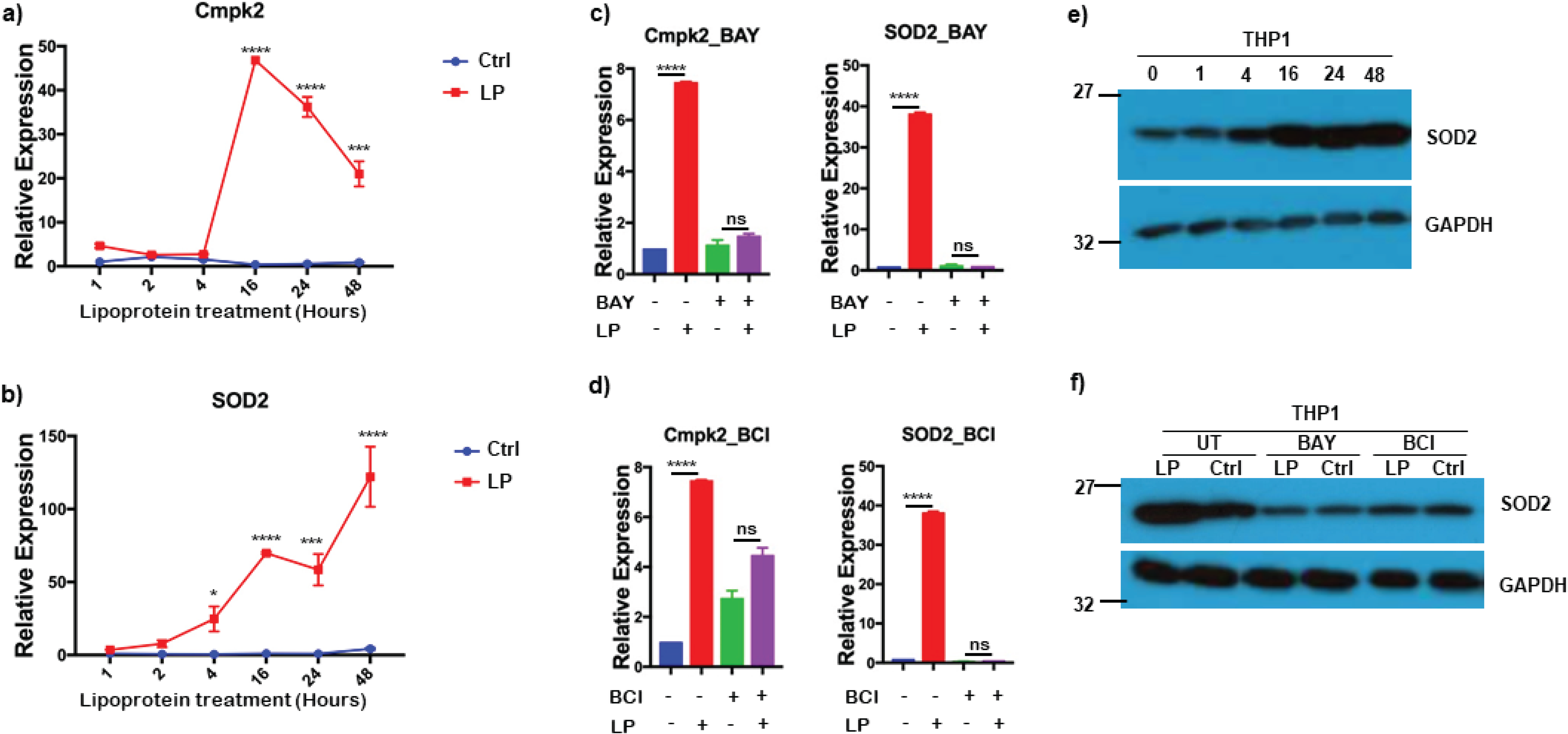
*Bb*LP upregulates mitochondrial stress related genes in THP1 cells. Line graphs showing the kinetics of relative expression of mitochondrial genes, a) *Cmpk2* and b) *Sod2* in LP-treated THP1 cells at 1,2,4,16,24 and 48 hours, determined by qPCR, while untreated THP1 cells were maintained as control. Bar graphs showing the *Bb*LP-induced relative expression of *Cmpk2* and *Sod2* genes in THP1 cells pre-treated with c) BAY and d) BCI inhibitors at 24 hours, determined by qPCR. Western blotting showing the expression of SOD2 protein in *Bb*LP-treated THP1 e) at 0,1,4,16,24 and 48 hours and f) pre-treated with BAY and BCI inhibitors, while untreated THP1 cells were maintained as control and GAPDH was used as normalization control. Data are presented as mean ± standard error of the mean (SEM). Statistical significance between samples is indicated as follows: *p < 0.05, **p < 0.01, ***p < 0.001, ****p < 0.0001.

Addition of both IKK inhibitor BAY and DUSP1 inhibitor BCI to PMA differentiated THP1 cells prior to treatment with *Bb*LP reduced the expression of *Cmpk2* and *Sod2* genes suggesting the role of NF-κB and DUSP1 mediated effects on mitochondrial oxidative stress responsive genes, *Cmpk2* and *Sod2* (**Fig. 13c and 13d**). Furthermore, immunoblot analysis of total lysates of PMA differentiated THP1 cells treated with *Bb*LP resulted in increased levels of *Sod2* at 4, 16, 24 and 48 hrs compared to levels at 0 and 1hpt (**Fig. 13e**). However, treatment with BAY or BCI prior to *Bb*LP addition reduced the levels of *Sod2* in differentiated THP1 cells indicating the modulatory effects of both IKK and DUSP1 on mitochondrial genes (**Fig. 13f**).

### *Bb*LP inhibits mitochondrial oxidative stress via MyD88 dependent pathway

While borrelial lipoproteins are known to signal through TLR2/1 and MyD88, we used MyD88-KO THP1 cells, differentiated into macrophages with PMA, to investigate the activation of mitochondrial oxidative stress-related genes by *Bb*LP at 24 hpt. Both *Cmpk2* and *Sod2* expression levels were reduced in MyD88-KO THP1 cells compared to parental THP1 cells (**Fig. 14a, b**). SOD2, a key antioxidant, plays an essential role in protecting cells from oxidative stress.

**Figure 14:**
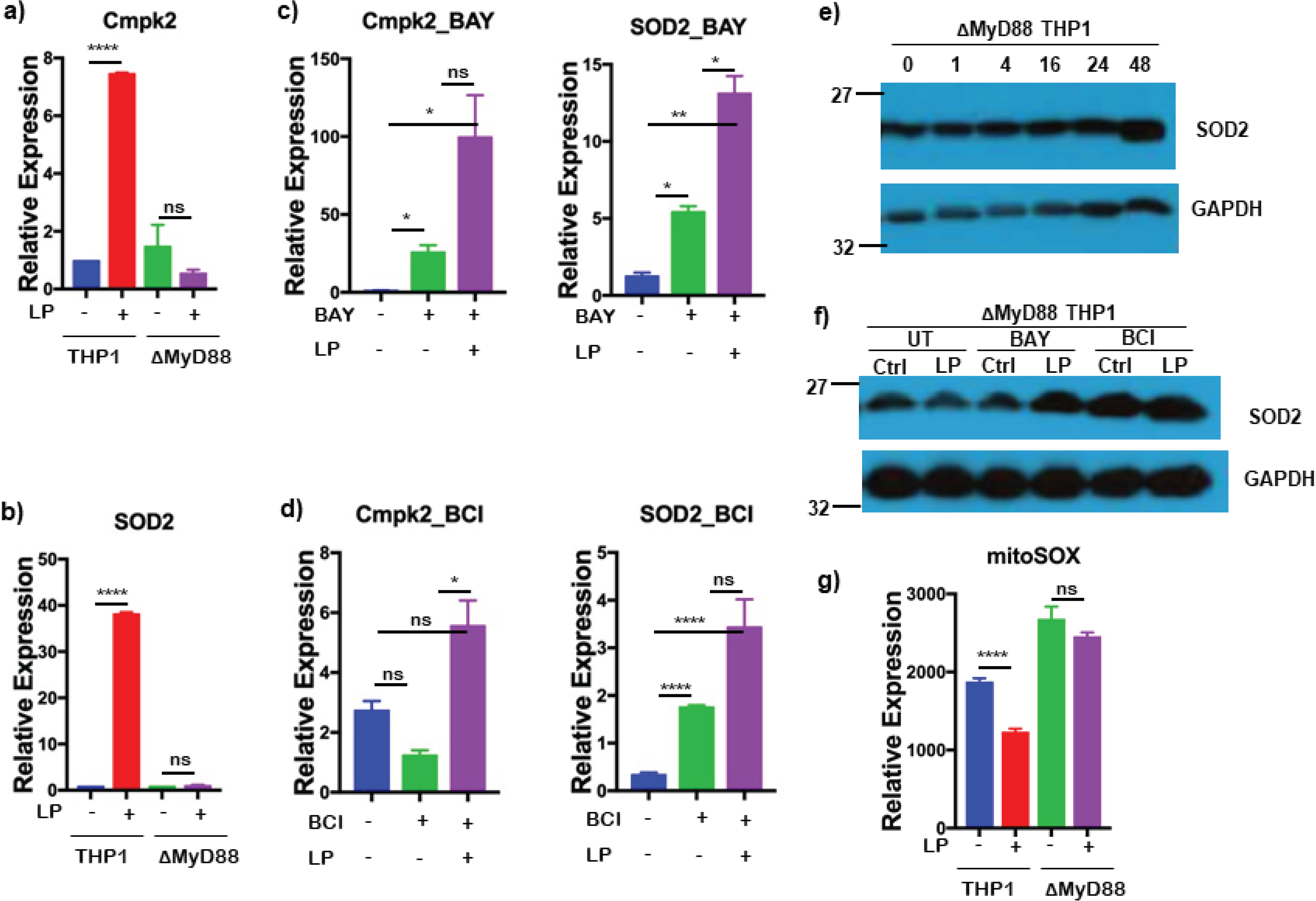
*Bb*LP-induced upregulation of mitochondrial stress is regulated by MyD88. Bar graphs showing the relative expression of mitochondrial genes, a) *Cmpk2* and b) *Sod2* in LP-treated MyD88 KO THP1 cells at 24 hours, determined by qPCR, while untreated cells were maintained as control. Bar graphs showing the LP-induced relative expression of *Cmpk2* and *Sod2* genes in MyD88 KO THP1 cells pre-treated with c) BAY and d) BCI inhibitors at 24 hours, determined by qPCR. Western blotting showing the expression of SOD2 protein in LP-treated MyD88 KO THP1 e) at 0,1,4,16,24 and 48 hours and f) pre-treated with BAY and BCI inhibitors, while untreated THP1 cells were maintained as control and GAPDH was used as normalization control. g) Bar graph showing the median fluorescence intensity of mitoSOX in THP1 and MyD88 KO THP1 cells with and without *Bb*LP treatment. Data are presented as mean ± standard error of the mean (SEM). Statistical significance between samples is indicated as follows: *p < 0.05, **p < 0.01, ***p < 0.001, ****p < 0.0001.

Treatment of MyD88-KO cells with BAY or BCI led to increased transcriptional levels of *Cmpk2* and *Sod2*, even without *Bb*LP, indicating the involvement of alternative pathways in the absence of MyD88 (**Fig. 14c, d**). No significant changes in SOD2 protein levels were observed in MyD88-KO cells, except at 48 hrs, which correlated with transcriptional data (**Fig. 14e**). SOD2 protein levels were elevated in BAY- or BCI-treated cells (**Fig. 14f**), similar to the transcriptional patterns of *Cmpk2* and *Sod2* in BbLP-treated MyD88-KO cells. These results suggest that inhibition of IKK and DUSP1 impacts mitochondrial stress regulation in macrophages in a MyD88-dependent manner.

In addition, we tested cellular ROS production using mitoSOX in differentiated macrophages. The mitoSOX assay measures oxidative stress by detecting superoxide production in mitochondria. The median fluorescence intensity of mitoSOX^hi^ cells was significantly reduced in the *Bb*LP-treated parental THP1 macrophages, while no difference in mitoSOX was observed in MyD88 KO cells, compared to the untreated controls (**Fig. 14g**). This suggests that *Bb*LP reduces mitochondrial ROS production in macrophages via a MyD88-dependent pathway, regulated by *Cmpk2* and *Sod2*, likely via their antioxidant properties.

## Discussion

The ability of spirochetes to modulate the host’s innate immune responses by exploiting components of tick saliva and its own PAMPs, such as lipoproteins, nucleic acids, peptidoglycan, and flagellin, plays a critical role in its survival during both the tick and mammalian phases of infection [11, 37, 42, 68–71]. The interactions between *Bb* and the host largely depend on the spatial and temporal expression of various lipoproteins on the surface, which engage with TLRs and non-TLR receptors, influencing *Bb*’s survival, colonization, and pathogenicity [72, 73]. Recently, scRNA-Seq analysis of murine splenocytes infected *ex vivo* with gfp-labelled *Bb* revealed induction of Dual specificity phosphatase 1 (*Dusp1*) and proinflammatory cytokines CXCL2 and IL1β in infected neutrophils compared to bystander neutrophils. Additional validation of scRNA-Seq data also showed a connection between increased levels of DUSP1, inflammatory cytokines and Caspase-3 levels in infected neutrophils compared to uninfected cells [59]. We extended these studies and performed scRNA-Seq with murine bone marrow-derived macrophages (BMDMs) exposed to purified borrelial lipoproteins for 1 and 4 hrs and compared the transcriptome changes with untreated BMDMs. Transcriptomic changes of individual DEGs, pathways and proteins were validated using qRT-PCR, proteomic, flow cytometry and specific kinase/phosphatase inhibitors with a focus on signaling events mediated by *Bb*LP integrating the role of DUSP1 in impacting proinflammatory, anti-inflammatory and mitochondrial responses (summarized in **Fig. 15**).

**Figure 15:**
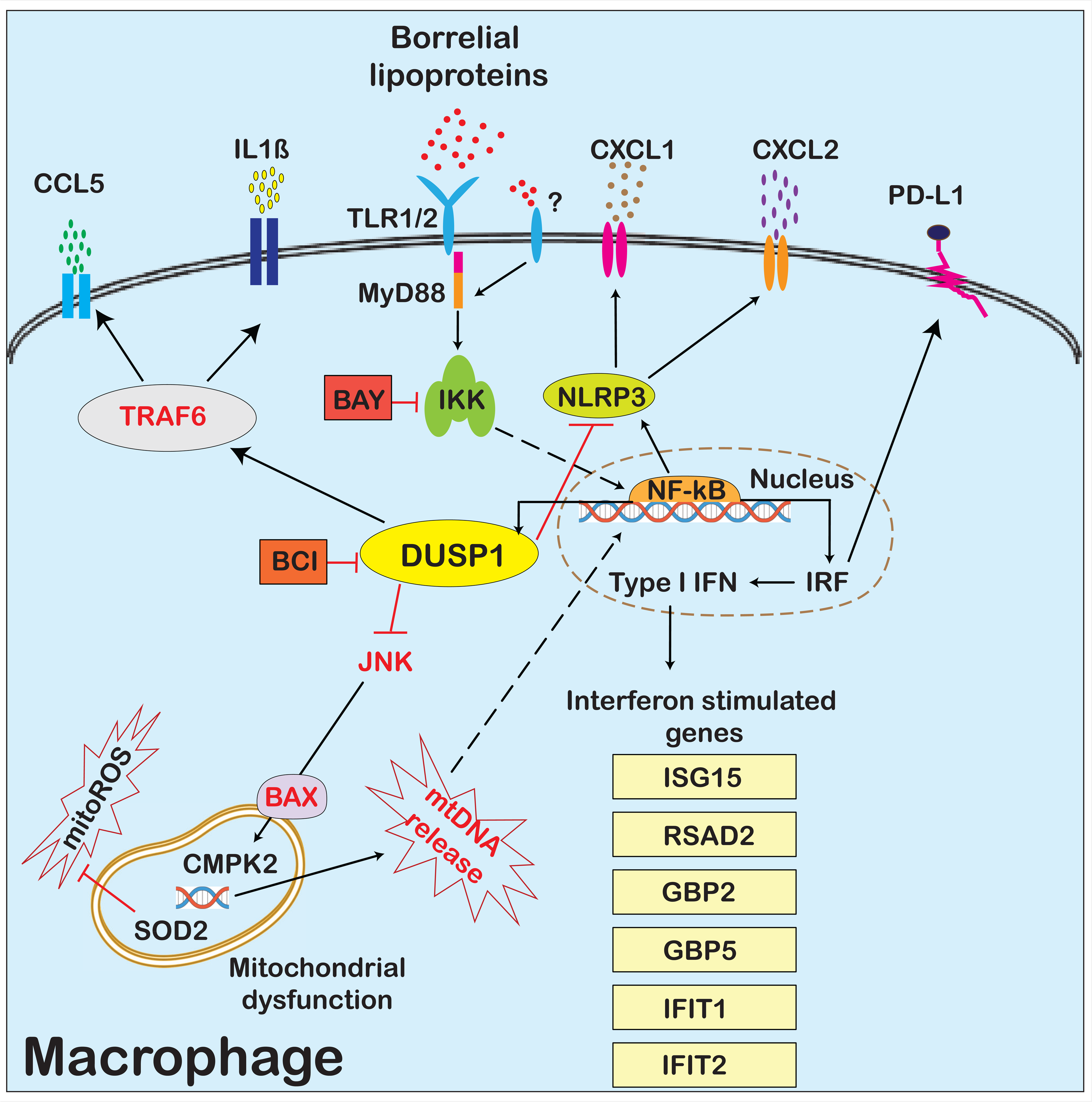
Overview of *Bb*LP induced changes in BMDMs. A graphical overview depicting the modulation of BMDMs by *Bb*LP. TLR1/2, bound to MyD88, are shown as the primary receptors for *Bb*LP. IKK and DUSP1 are highlighted as central regulators, marked in green and yellow, respectively, while their inhibitors BAY and BCI are shown in red boxes. Mitochondrial effectors like mitoROS and mtDNA are depicted as stars. Type-I-IFN-regulated genes are listed in yellow boxes. Solid arrows indicate gene regulation pathways, while dotted arrows represent the translocation of proteins into the nucleus. Cytokines such as CXCL1, CXCL2, CCL5, and IL1B are shown as various colored granules in the extracellular space. The previously identified mediators, including JNK, BAX, TRAF6, and mtDNA, which were not experimentally examined in this manuscript, are highlighted in red.

A heterogeneous population of BMDMs, consisting of 11 distinct clusters, was identified across all three samples (untreated, LP1, LP4) based on DEGs (Fig. 1a, b, and c). However, all clusters expressed the leukocyte marker *Ptprc* (CD45) and the macrophage marker *Csf1r*, confirming that these clusters represent macrophage subtypes (Fig. 1c, d). Furthermore, macrophage heterogeneity was evident based on DEGs within the samples, defining three functional classes of BMDMs: 1) inflammatory macrophages (*Fgf13*, *Ccnd1*) [74]; 2) antigen-presenting macrophages (*Apoe*, *Slc9a9*) [75]; and 3) cell cycle-related macrophages (*Hist1h1b*, *Hist1h2ap*) [76], identified based on the functionally relevant genes (**Fig. 1d, 1e**). Inflammatory macrophages constitute the major population exhibiting significant DEGs between untreated and lipoprotein-treated BMDMs with several hub genes associated with pro-inflammatory functions. This is consistent with other scRNA-Seq studies that have identified inflammatory macrophages with similar sets of hub genes in various inflammatory disease conditions [77, 78].

In the Harmony-integrated UMAP plots of the control, LP1, and LP4 samples, the LP4 clusters are distinctly separated, while BMDM subsets remained consistent between untreated (Ctrl) and lipoprotein-treated (LP1) samples (**Fig. 2a,b**). These findings were further supported by pseudotime analysis (**Supp-Fig. 3**). Pseudotime analysis of atherosclerotic macrophages showed that several macrophage subsets transitioned into M1 macrophages, aligning with multiple genes identified in this study reflecting broader responses of macrophages to activating signals [79]. Many upregulated genes in the LP4 clusters were specific to classical M1 polarization (**Supp-Fig. 1**), including *Nos2*, *Cd40*, and *Ch25h* [80, 81]. These results suggest that lipoproteins drive M1 polarization in macrophages, which are known to produce pro-inflammatory cytokines through transcription factors such as STAT1, NF-κB, and IRF-5, [82], along with additional macrophage phenotypes broadening the innate immune responses following *Bb* exposure (**Supp-Table 1**). Moreover, receptors such as *Clec4d* and *Clec4e*, which were differentially regulated following *Bb* treatment, correlate the effects observed with purified lipoproteins [50, 83, 84]. Importantly, pathways regulated by growth factor receptors, including phosphatidylinositol 3-kinase pathways (*Rab7b, Pik2, Pik3cb*) and mitogen-activated protein kinase/extracellular signal-regulated kinase (MAPK/ERK) pathways (*Dusp1*, *Dusp4*, *Dusp5*), as well as downstream components such as AP-1 transcription factors (*Fos, Fosb*) and early growth response genes (*Egr1, Egr2*) [85] and mitochondrial energy metabolism genes (*Ndufv3*, *Ndufa2*, *Uqcr10*, *Uqcr11*, *Atp5j2*) were down-regulated in lipoprotein treated BMDMs (**Fig. 2c**). Down-regulation of these signatures suggest that lipoproteins suppress mitochondrial energy metabolism and inhibit the growth response, leading to impaired immune function in macrophages, where dual specificity phosphatases play a key role in regulating the MAPK and PI3K pathways [86–88]. This suggests that lipoproteins modulate multiple components in macrophages, impacting immune response efficiency associated with elevated inflammation during spirochete infection [89].

Among the identified DEGs, macrophage surface receptors such as *Tlr2*, *Myd88*, and *Il13ra1*, along with ISGs like *Gbp2*, *Gbp5*, *Ifit1*, *Ifit2*, and *Isg15*, were expressed across all subsets of BMDMs (Supp-Fig. 2), consistent with previous findings [39, 45, 67]. The upregulation of these receptors and ISGs in inflammatory macrophages in response to *Bb*LP correlates observations in macrophages from atherosclerosis models [90]. In addition to classical receptors like TLR2/MyD88, receptors such as *Cd274* (PD-L1), *Nr3c1*, and *Ldlr* were upregulated in LP4, suggesting novel roles for these receptors in macrophage interactions with lipoproteins. PD-L1 expressing macrophages are more mature/activated and promote CD8^+^ T cells proliferation and cytotoxic capacity in tumor environment [91]. *Nr3c1*, a glucocorticoid receptor that is epigenetically regulated, and *Ldlr*, a low-density lipoprotein receptor involved in lipid metabolism, both play crucial roles in controlling inflammation in macrophages, highlighting a complex and multi-faceted macrophage response [92, 93].

Analysis and validation of the scRNA-Seq data sets between LP1 and LP4 in comparison with untreated BMDMs has been directed at three broad areas of focus namely 1) significance of mitochondrial and ribosomal genes affecting the metabolic status of the BMDM; 2) effects of DUSP1 upregulated in LP1 and 3) impact of ISGs and other signaling genes notably chemokines (**Fig 3 a, b and c**). There also appears to be a unique subcellular localization profile with DEGs upregulated in the untreated BMDMs localized to the mitochondria, while those in LP1 and LP4 localized to endosomal compartments/lysosomes and Golgi apparatus, respectively (**Fig. 3d, e and f**). These observations add to the significance of the breakdown products of *Bb*, an extracellular pathogen, in modulating cellular processes influencing macrophage responses and cumulative innate immune responses induced in a mammalian host.

Proteomic analysis of the total lysate from *Bb*LP-infected BMDMs validated levels of DEGs at the protein level between control, LP1 and LP4 samples (**Fig. 4**). Interleukin receptor, IL13RA1 was upregulated in LP4 clusters at both the transcriptional (**Supp-Fig. 2**) and protein levels (**Fig. 4f**). However, genes encoding the *Il13ra2* subunit, as well as *Il13* and *Il4* genes, were not expressed in untreated BMDMs. The IL13RA1 subunit forms a receptor complex with IL4RA, serving as the primary receptor for IL13 and IL4 [94]. Binding of IL13 and IL4 proteins to tyrosine kinase TYK2 may mediate signaling processes leading to the activation of JAK1, STAT3, and STAT6, resulting in interferon signaling [95]. The presence of peptides specific to MyD88, GBP5 and PD-L1 reflected the transcriptomes upregulated in LP4 compared to LP1 or untreated BMDMs validating the scRNA-Seq findings (**Fig. 4d**). The significance of IL13RA1, however, remains to be investigated although it was detected in mouse blood in response to *Bb* [96]. Stimulation of toll-like receptors (TLRs) by pathogens activates signaling cascades, leading to the translocation of nuclear factor-κB (NF-κB) to the nucleus [97] and the activation of interferon regulatory factors 3/7 (IRF3/7) [98]. Consistent with these known mechanisms, both scRNA-Seq (Fig. 3 b,c) and proteomic analyses (**Fig. 4e**) revealed MAPK phosphorylation-mediated NF-κB and IRF pathways in BMDMs exposed to *Bb*LP (**Fig. 5,9,12**).

In addition, unbiased proteomic analysis revealed 21 matches to *B. burgdorferi* proteome with peptide abundance for lipoproteins such as outer surface proteins A, B, C, D and 6.6 kDa lipoprotein. Since the lysates were prepared from macrophages, after washing with PBS, only internalized borrelial proteins were preferentially detected, suggesting that these proteins were processed at higher levels in macrophages. Even though we did not observe any significant levels of the flagellin (FlaB) in the detergent phase by immunoblot assay (**Supp-Fig. 1**), the presence of peptides specific to flagellin indicate either the sensitivity of method used (nanoLC-MS/MS) or the potential for association of FlaA/FlaB with other abundant lipoproteins (**Fig. 4c**). However, the abundance of lipoproteins likely induced significant macrophage activation reflected in the DEGs in various clusters of macrophages analyzed in this study [99].

While several validations of the scRNA-Seq data either confirmed previous findings from bulk RNA sequencing analyses regarding responses to *Bb*LP [84, 100] or identified novel DEGs involved in enhancing or suppressing the innate immune responses mediated by macrophages in response to *Bb*LP, we focused more closely on the key DEG, *Dusp1*. A major finding of this study is the transient expression of DUSP1 in LP1 compared to LP4, as shown in the scRNA-Seq analysis (**Fig. 2d**), which was further validated at both the transcriptional and translational levels (**Fig. 5a, b**). DUSP1 dephosphorylates threonine and tyrosine residues on mitogen-activated protein kinases (MAPKs), preferentially p38 and c-Jun N-terminal kinase. By rendering these kinases inactive, DUSP1 serves as an excellent drug target for the positive and negative regulation of immune responses [101]. However, the significance of DUSP1 in modulating the innate response induced by borrelial lipoproteins is unclear. While there was a significant increase both at the transcriptional and translational level at 1 hr of treatment with *Bb*LP, there was a rapid decline at the 4hr time point suggesting rapid post-translational decay (**Fig. 5a, b**). This pattern is consistent with the response observed in mouse spleen following LPS injection, where *Dusp1* gene expression was high at early hours but low at 24 hpt [102]. Moreover, in our previous study, we demonstrated that intact *Bb* induces *Dusp1* upregulation in splenic neutrophils and macrophages, highlighting its role in modulating the CXCL1 and CXCL2 axis in neutrophils during *Bb* infection and underscoring the role of DUSP1 in early stages of innate immune response [59].

The role of DUSP1 in the signaling events mediated by *Bb*LP via TLRs, MyD88 and IKK were further interrogated using specific inhibitors targeting either IKK or DUSP1. BAY 11-7082, an inhibitor of IKK, prevented the upregulation of DUSP1 by lipoproteins, suggesting a key regulatory role for IKK in regulating the NF-κB resulting in increased levels of DUSP1 in macrophages and thereby influencing early modulation of inflammatory responses (**Fig. 5c**). As shown in **Fig. 5c** and **5d**, pre-treatment of BMDMs with 2-benzylidene-3-(cyclohexylamino)-1-indanone hydrochloride (BCI), a specific allosteric inhibitor of DUSP1 and DUSP6, before *Bb*LP stimulation, completely inhibited DUSP1 expression, confirming that BCI effectively suppresses DUSP1 protein expression in macrophages [103].

A significant increase in the expression of NLRP3-associated genes such as *Nlrp3*, *Il1f9*, and *Ccrl2*, independent of the *Bb*LP dose, was observed in treated BMDMs although the levels of *Nlrp3* was significantly lower at the 24hr time point consistent with prior observations (**Fig. 6a**) [104, 105]. Both the transcriptional and protein levels of CXCL1 and CXCL2 were elevated in macrophages exposed to *Bb*LP (**Fig. 6a, b**). Interestingly, treatment with the BCI inhibitor significantly enhanced the transcriptional and cytokine levels suggesting that DUSP1 may act as a negative regulator of the NLRP3 inflammasome-mediated CXCL1 and CXCL2 expression in macrophages exposed to lipoproteins (**Fig. 6c, 8a**). This finding aligns with previous research demonstrating enhanced CXCL1 and CXCL2 activation in DUSP1-deficient macrophages following lipopolysaccharide (LPS) stimulation although the TLR involved in LPS driven signaling events is different [106]. Inhibition of IKK with BAY, however, resulted in increased levels of transcripts specific to *Cxcl1* and *Cxcl2* (**Fig. 6c**) although the level of the cytokines detected in the supernatants were at the same level as from untreated cells indicating a positive regulatory role for IKK in NLRP3 inflammasome-mediated CXCL1 and CXCL2 expression in macrophages (**Fig. 8a**). The NF-κB and STAT1 pathways are well-established regulators of CXCL1 and CXCL2 transcription [107]. The CXCL1→CXCR2 axis is critical in cellular senescence and immune cell recognition, underscoring its importance in inflammatory responses mediated by macrophages [108, 109]. Inhibitors of IKK-mediated NLRP3 responses notably the dampening of the release of inflammatory cytokines such as CXCL1 and CXCL2 are likely candidate formulations to minimize the pathological effects of *Bb* or its PAMPs[110].

It has been shown that DUSP1 mediates TRAF6 dependent regulation of levels of CCL5 and IL1B [111, 112]. While *Bb*LP upregulated *Ccl5* and *Il1b* genes and their cytokine levels in BMDMs, with peak gene expression observed at 24 hpt (**Fig. 7a**), treatment with both IKK and DUSP1/6 inhibitors significantly inhibited the transcriptional and cytokine levels of CCL5 and IL1B compared to untreated *Bb*LP-infected macrophages (**Fig. 7b, 8a**), indicating the essential roles of IKK kinases and DUSP1 in mediating TRAF6 dependent induction of NLRP3-independent proinflammatory cytokines such as IL1B [105]. The relevance of these findings is underscored by recent studies demonstrating the therapeutic potential of targeting CCL5 and IL1B in various inflammatory conditions such as in restoration of immune homeostasis and reduction in viral load in critical COVID-19 patients through disruption of the CCL5/RANTES-CCR5 pathway [113]. Similarly, blocking IL1B activity has shown rapid and sustained reduction in disease severity in autoinflammatory syndromes [114]. Furthermore, IL1B-expressing macrophages have been implicated in inflammatory arthritis interacting with CD8+ T cells via the CCL5/CCL3 axis, serving as potential therapeutic targets for ameliorating a range of inflammatory conditions [115]. Using a well-established TLR2 agonist and PAMP associated with *Bb*LP - Pam3CSK4, it was possible to demonstrate that *Cxcl1*, *Cxcl2*, *Ccl5* and *Il1b* were upregulated in BMDMs. However, treatment with either BAY or BC1 did not reduce the cytokine levels of CXCL1 and CXCL2 although the levels of CCL5 and IL1b were lower in the presence of the inhibitors. One possible explanation for this discrepancy is levels of engagement of TLRs by the PAMP alone versus PAMP being a part of a larger sets of proteins as would be expected with *Bb*LP.

*B. burgdorferi* elicits robust innate and inflammatory responses involving both IFN-I (IFN-α and β) and IFN-II (IFN-γ) cytokines that drive the expression of interferon-stimulated genes (ISGs) [116, 117]. *Bb*LP upregulate several ISGs, including *Gbp2*, *Gbp5*, *Ifit1*, *Isg15*, *Rsad2*, and *Hcar2*, showing an increasing trend over the course of infection (**Fig 9a**). These observations are consistent with previous study demonstrating the significance of IFN pathway-related genes, such as *Ifnb*, *Cxcl10*, *Gbp1*, *Ifit1*, *Ifit3*, *Irf7*, *Mx1*, and *Stat2*, that were differentially expressed in mice infected with *B. burgdorferi* [96]. While treatment with IKK inhibitor BAY reduced the levels of *Gbp2*, *Gbp5*, *Ifit1*, *Isg15*, *Rsad2*, and *Hcar2* significantly, the levels of reduction in response to DUSP1/6 inhibitor BCI only partially reduces ISG expression, suggesting a secondary role for DUSP1/6 in this context (**Fig. 9b**). This implies that targeting IKK kinases could potentially impair the host’s ability to clear *B. burgdorferi,* whereas targeting DUSP1/6 may offer a more nuanced approach to modulating the immune response against *Bb* infection in reservoir hosts [43]. Among ISGs, guanylate-binding proteins (GBPs) have been shown to contribute to host defense against pathogens and to promote inflammasome activation during bacterial infections [118]. GBP2 is elevated in various tissues during *Bb* infection in mice, underscoring its involvement in the innate immune responses [5]. Both GBP2 and GBP5 exhibit antiviral activities and play roles in immune responses, with GBP5 specifically promoting NLRP3 inflammasome assembly [119, 120]. Moreover, treatment with an IKK inhibitor completely inhibits the expression of these ISGs in *Bb*LP-infected BMDMs, emphasizing the critical role of IKK kinases in their regulation (**Fig. 9b**).

The study of programmed cell death protein 1 (PD-1) and its ligand PD-L1 has significant implications in immunotherapy, particularly in diseases like Lyme arthritis [121]. Recently, upregulation of PD-1 on CD4+ T cells and PD-L1 on antigen-presenting cells was demonstrated in Lyme arthritis suggesting a role for the PD-1/PD-L1 pathway in modulating T cell responses and inflammation induced by *Bb* [121]. Both scRNA-Seq and proteomic analyses revealed that PD-L1 encoded by *Cd274* is markedly expressed in macrophages in response to *Bb*LP in LP4 sample (**Fig. 4d**). This was further confirmed using flow cytometry and antibodies specific to PD-L1 on macrophages following treatment with *Bb*LP (**Fig. 10 a-c**). Inhibition of DUSP1/6 with BCI completely blocked PD-L1 expression in macrophages prior to *Bb*LP treatment (**Fig. 10 d-f**). Peak levels of PDL-1 was observed at 24hrs by immunoblot analysis and corroborated using confocal microscopy showing PDL-1 on the surface of BMDMs at 24 hrs, which was inhibited by the DUSP1/6 inhibitor BCI (**Fig. 11 a,c**). PD-L1 was also upregulated in BMDMs in response to Pam3CSK4 similar to *Bb*LP (**Fig. 11b**) These observations aligns with previous studies showing that DUSP1 can regulate PD-L1 expression in the context of microbial infections potentially enhancing the accumulation of PD-1+CD4+ T cells and exacerbating inflammation in response to *Bb* infection [122, 123]. Notably, targeting the PD-1/PD-L1 pathway did not impact *Bb* clearance directly although it affected T cell expansion in mice [121]. Recent studies demonstrated that PD-L1 expressing macrophages are hyperactive, mature and immunostimulatory and plays a key role in clearing tumors [91]. Therefore, targeting DUSP1 to modulate PD-L1 expression might be a novel strategy to improve the clearance of *Bb* from the site of infection.

*Bb*LP upregulated NF-κB-regulated Nlrp3-associated genes (**Fig. 6**) and induced IRF-regulated expression of ISGs in murine BMDMs via activation of IKK (**Fig. 9**). Similarly, *Bb*LP treatment elicited both SEAP and Luciferase activities in THP1-Dual reporter cells, indicating activation of both NF-κB and ISGs in human macrophages (**Fig. 12 a**), consistent with the response observed in murine BMDMs. SEAP and Luciferase activities were significantly lower in MyD88-KO-THP1 cells compared to THP1 reporter cells during *Bb*LP treatment (**Fig. 1**2 b, c), demonstrating that MyD88 is essential for the activation of these pathways in macrophages. Treatment with BAY and BCI inhibitors in both THP1 cells and MyD88-KO cells significantly reduced SEAP and Luciferase activities induced by *Bb*LP treatment highlighting the requirement of IκB kinases and DUSP1/6 phosphatases for NF-κB and ISG activation in both human and murine macrophages (**Fig. 12d-f**). Prior studies have demonstrated that BCI (DUSP1/6 inhibitor) attenuates lipopolysaccharide-induced inflammatory responses in murine macrophage cells by inhibiting the NF-κB pathway [124]. These observagions further support that BCI inhibits NF-κB and ISG-mediated inflammatory responses in both murine and human macrophages. SEAP activity in BCI-treated (20µM) THP1 cells was significantly increased during Pam3CSK4 stimulation, whereas it was inhibited during *Bb*LP stimulation (**Fig. 12 g-i**). These results are indicative of *Bb*LP activation of NF-κB and ISGs in macrophages is dependent on TLR2/MyD88, while DUSP1’s involvement in NF-κB and ISG activation is independent of the TLR2/MyD88 cascade, consistent with findings in murine BMDMs [125].

scRNA-Seq revealed downregulation of several mitochondrial genes related to ROS production following treatment of BMDMs with *Bb*LP and upregulation of mitochondrial antioxidant genes such as cytidine monophosphate kinase 2 (*Cmpk2*) and mitochondrial superoxide dismutase 2 *Sod2* (**Fig. 3 a,b**). These observations were validated in THP1 macrophages using qPCR (**Fig 13 a,b**), suggesting reduced ROS production. The transcriptional levels of *Cmpk2* peaked at 16 hpt, aligning with previous studies peak levels were observed at 12 hpt following LPS treatment in macrophages [126]. CMPK2, is an early and highly expressed ISG in macrophages following bacterial and viral infections, as well as in response to lipopolysaccharide (LPS) and polyinosinic:polycytidylic acid (poly IC) [126–128]. SOD2 plays a crucial role in developing oxidative stress resistance in M1 macrophages [129]. Treatment with *Bb*LP resulted in increased transcriptional and protein levels of SOD2 in THP1 macrophages (**Fig. 13 b,c**) while no such effect was observed in MyD88 knockout (KO) cells (**Fig. 14 a,b**) indicating that mitochondrial stress response is partly regulated by MyD88. Moreover, the mitoSOX assay showed that *Bb*LP significantly reduced cellular ROS activity in THP1 but not in MyD88 KO cells, indicating that *Bb*LP reduces mitochondrial ROS (mitoROS) in a MyD88-dependent manner (**Fig. 14g**). *Bb* decreases the long-term capacity of PBMCs to generate ROS and evade killing while modulating the host immune system [130], consistent with our findings. A recent study demonstrated that borrelial outer membrane vesicles modulate mitochondrial oxidative stress in human neuroblastoma cells [131]. These observations suggest that *Bb*LP treatment inhibits ROS production in macrophages, regulated by SOD2 and CMPK2 in a MyD88-dependent manner.

BAY and BCI inhibited the transcriptional expression of *Bb*LP-induced *Cmpk2* (**Fig. 13e**) and *Sod2* in THP1 macrophages (**Fig. 13f,g**). Similarly, treatment with BAY or BCI prior to *Bb*LP addition reduced the protein levels of SOD2 in differentiated THP1 cells, suggesting that both IKK and DUSP1 regulate mitochondrial stress in macrophages in response to borrelial lipoproteins. Regulation of SOD2 by the NF-κB pathway has been identified during epithelial-mesenchymal transition, and SOD2 induction in transformed keratinocytes was concurrent with the suppression of TGF-β-mediated induction of both ROS and senescence [132]. However, BAY and BCI inhibitors elevated the expression of *Cmpk2* and *Sod2* in MyD88-KO cells, independent of lipoprotein treatment (**Fig. 14c-e**). Previous studies have shown that NF-κB and DUSP1 modulate mitochondrial transcription factors and ROS suggesting that inhibition of NF-κB and DUSP1 affects mitochondrial stress homeostasis in macrophages in a MyD88-dependent manner [128, 133, 134]. Moreover, LPS stimulation in BMDM increases ROS, whereas *Bb*LP reduces ROS, indicating that the *Bb*LP-induced response is distinct from the LPS-stimulated response [135].

Mitochondrial dysfunction is observed in *Bb*LP-treated macrophages, as indicated by the downregulation of mitochondrial energy metabolism and protein synthesis pathways (Fig. 3d), associated with reduced ROS (**Fig. 14g**). This dysfunction likely impairs efferocytosis, leading to an accumulation of dead cells and debris [136]. In turn, several pro-inflammatory signaling cascades were activated (**Fig. 7,8**). The resulting increase in inflammatory response likely contributes to immunosuppression, compromising the overall immune function [137]. Despite this mitochondrial dysfunction, macrophages exhibit active phagocytosis of *Bb*LP at an early stage, as evidenced by the upregulation of endosomal genes at 1 hpt (**Fig. 3e**) [138]. This indicates that macrophages are still capable of recognizing and internalizing *Bb*LP. Further, at 4 hpt, the upregulation of lysosome- and vesicle-related genes suggests mechanisms of processing and degradation of internalized lipoproteins by macrophages (**Fig. 3f**). These sequential modulations underscore the ability of macrophages to effectively process borrelial lipoproteins, despite the mitochondrial challenges. Further studies are required to understand the relevance of these mechanisms in immune suppression in macrophages by *Bb*LP.

## Conclusion

In conclusion, this study elucidates the diverse responses of macrophages to borrelial lipoproteins. We identified robust inflammatory responses characterized by elevated expression of CXCL1, CXCL2, IL1B, CCL5 and other key cytokines and chemokines, regulated through NF-κB and ISG pathways, in response to *Bb*LP. Our findings highlight the critical role of IFN-stimulated genes (ISGs), including *Gbp2*, *Gbp5*, *Ifit1*, *Isg15*, *Rsad2*, and *Hcar2* as well as PD-L1 in supporting host defenses against *Bb* infection. Crucially, blocking IKK kinases and DUSP1 critically modulates the *Bb*LP-induced cytokines, type-I IFN and mitochondrial response in macrophages (summarized in **Fig. 15**). These insights into *Bb*LP-induced immune activation provide a platform for targeted therapies aimed at modulating immune responses in Lyme disease and related infections, warranting further investigation in animal models. Single cell RNA-Seq data provides a novel list of genes involved in the activation of inflammatory macrophage subsets by borrelial lipoproteins, advancing our understanding of the pro-inflammatory subset of macrophages and their responses to infection with the agent of Lyme disease.

## Materials and Methods

### Mice and Ethics statement

All animal experiments were conducted following NIH guidelines for housing and care of laboratory animals and in accordance with protocols approved by the Institutional Animal Care and Use Committee (protocol number MU071) of The University of Texas at San Antonio (UTSA). Based on these guidelines, general condition and behavior of the animals were monitored by trained staff or by laboratory personnel. The animal facilities at UTSA are part of Laboratory Animal Resources Center (LARC), which is an AAALAC International Accredited Unit. Six-week old female C3H/HeN mice (Charles River Laboratories, Wilmington, MA) was used in this study.

### Bacterial strain and growth conditions

A low passage infectious clonal isolate of *Borrelia burgdorferi* B31-A3 were propagated in liquid Barbour-Stoenner-Kelly (BSK II) media supplemented with 6% normal rabbit serum (Pel-Freez Biologicals, Rogers, AR) with appropriate antibiotics (Sigma-Aldrich, St. Louis, MO) as previously described [139–144]. Bacteria were grown under conditions that induced lipoproteins and when the cultures reached a density of 1×10^7^ spirochetes/ml, the count and viability of spirochetes was confirmed before lipoprotein extraction.

### Isolation of lipoproteins from *B. burgdorferi*

Lipoproteins were extracted from wildtype *B. burgdorferi* B31-A3 strain as described previously [145]. Briefly, 1x10^9^ *Bb* was resuspended in 1 mL of PBS (Phosphate Buffered Saline) pH7.4 containing 1% Triton X-114 (Sigma) by gently rocking at 4°C overnight. The TX-114 insoluble material was removed by two centrifugations at 15,000xg at 4°C for 15 mins. The supernatant was transferred to a sterile tube and incubated at 37°C for 15 mins. Then the mixture was centrifuged at 15,000xg for 15 mins at room temperature (RT). The top aqueous phase was transferred to a new tube and re-extracted one more time with 1% TX-114 as described above. The lower detergent phase was washed with 1 mL PBS pH 7.4 three times. Protein in the final detergent phase was precipitated by adding 10-fold volume of ice-cold acetone followed by centrifugation at 15,000xg at 4°C for 30 mins. Following vacuum drying, *B. burgdorferi* lipoproteins (*Bb*LP) pellets were resuspended in sterile 1X PBS and stored at -20°C until further use. Lipoprotein integrity was checked through Coomassie stained SDS-PAGE gel, and the lipoprotein concentration was quantified using BCA assay kit (Thermo Scientific).

### Bone marrow derived macrophages culture

Bone marrow cells were collected from femurs of normal female (*n=5*) C3H/HeN mice, pooled, and centrifuged at 500xg for 5 minutes at 4°C. The cell pellet was resuspended in 4 mL of 0.14M ammonium chloride solution to lyse RBC. After 5 minutes incubation at RT, RBC lysis was stopped by addition of 8 mL of fresh RPMI supplemented with 10% Fetal bovine serum (R10, FBS, Hyclone), centrifuged at 500xg for 5 minutes at 4°C. The bone marrow cells were resuspended in 40 mL of R10+15% L929 Cell Conditioned Media (LCCM) and passed through 70 µM cell strainer. In order to generate bone marrow derived macrophages, 10 mL of cell suspension at a density of 1x10^6^/mL was plated in petri dishes (CELLTREAT Scientific Products, MA, USA) and incubated at 37°C. Following replacement of 70% media with fresh R10 media after 4 days from each Petri dish, the cells were allowed to differentiate for 8 days at which time the culture media was removed and washed with 10 mL of cold PBS twice and treated with 2 mL of Accutase™ Cell Dissociation Reagent (Stem cell). After 20 minutes of treatment with Accutase at 37°C, the plates were tapped to detach the bone marrow derived macrophages (BMDMs) and were washed once with 4 mL of fresh R10 media and centrifuged at 500xg for 5 minutes. The BMDMs were resuspended and seeded at 1x10^6^ cells/mL in fresh R10 media without Pen/Strep in non-culture treated 6-well plates (Corning). After overnight incubation, the cells were treated with 1 mL of *Bb*LP (1, 0.1 or 0.01 µg/mL) resuspended in R10 media. PBS treated BMDM was maintained as control. After 1, 4 or 24 hpi, culture supernatant was removed, stored at -80°C for cytokine analysis. BMDMs were detached using Accutase, washed with cold PBS, checked for viability using Trypan Blue and processed for isolation of RNA or protein and for analysis using flow cytometry.

### THP1 and THP1-KO-MyD Dual reporter cells

THP1 and THP1-KO-MyD Dual reporter cells (Invivogen) were cultured in RPMI 1640, 2 mM L-glutamine, 25 mM HEPES, 10% heat-inactivated fetal bovine serum, 100 μg/ml Normocin™, Pen-Strep (100 U/ml-100 μg/ml) in T75 flasks. THP1 monocytes were differentiated to macrophages after 3-h exposure to 50 ng/ml phorbol 12-myristate 13-acetate (PMA, Sigma–Aldrich) in 1 mL of growth medium. Cells were washed with fresh medium to remove non-adherent cells. Cells were treated with 100 µL of growth medium containing BAY 11-7082 (IkappaB-IκB kinase inhibitor, Cat# B5556, Sigma) or *(E)*-2-Benzylidene-3-(cyclohexylamino)-2,3-dihydro-1H-inden-1-one (BCI, DUSP 1/6 inhibitor, Cat# 317496, Sigma) at different concentrations as indicated for each experimental condition. Plates were incubated at 37°C under 5% CO_2_ for 1 hr and were treated with 1 µg/mL of *Bb*LP or PBS. At 1, 4, 16 and 24 hours post-treatment (hpt) culture supernatants were collected to detect and quantify the Secreted embryonic alkaline phosphatase (SEAP) by measuring the absorbance at 655 nm or secreted luciferase using Varioskan luminometer (Thermo scientific), using Quanti-Blue and Quanti-Luc reagents as per manufacturer’s protocol [66].

### Treatment of macrophages with inhibitors

To understand the roles of IκΒ kinase complex and DUSP1/6 phosphatases in the regulation of *Bb*LP induced response in macrophages, BMDM were grown overnight in 6-well plate. Supernatant was removed, washed with PBS. BAY 11-7082 (IκB kinase inhibitor, Cat# B5556, Sigma) is an inhibitor of I IκBα phosphorylation and degradation, inhibits stimulant-induced IκBα phosphorylation with IC50 of 10 μM in tumor cells [146]. BCI ((2-benzylidene-3-(cyclohexylamino)-1-Indanone hydrochloride), Cat# 317496, Sigma) has been identified as a biologically active allosteric inhibitor of DUSP1 and DUSP6, serving as a DUSP1/6 specific inhibitor at 10 µM using zebrafish screening model [103]. To inhibit IKK and DUSP1/6 activity in macrophages, BMDM or PMA-treated THP1 cells were pre-treated with 1 mL of R10 containing 15 µM Bay 11-7082 or 10 µM BCI at 37°C in CO_2_ incubator for 1 hr, followed by treatment with 1 µg/mL of *Bb*LP or PBS for 1, 4 or 24 hpi. Culture supernatants were collected for cytokine analysis and cells were subjected to RT-PCR analysis to study the expression of the identified DEGs or for analysis by flow cytometry to determine expression levels of select markers in BMDMs.

### Single cell RNA: Library preparation and sequencing

Single-cell RNA-seq (scRNA-seq) analysis was performed to determine the cellular profiles and transcriptomes of individual cells following stimulation with borrelial lipoproteins. Briefly, 1x10^6^ BMDMs treated with 1µg/mL of *Bb*LP were collected at 1 and 4 hpt and PBS-treated BMDMs were collected at 4 hpt as described previously [59]. The single cell suspensions were subjected to droplet-based single cell RNA sequencing using Chromium Single Cell 3’ (v3.1) Reagent Kit as per manufacturer’s protocol and as reported previously [59]. We aimed to capture 5000 cells/lane and each cell was labeled with a specific barcode, and each transcript labeled with a unique molecular identifier. The library was generated following the manufacturer’s recommendations for the 3′ Gene Expression v3 kit, followed by Illumina sequencing of each library and FASTQ files were generated.

### Single cell RNA seq: Data processing

FASTQ files generated from the scRNA-Seq were uploaded to the 10x genomics cloud (https://cloud.10xgenomics.com/cloud-analysis) and the data was processed using the Cell Ranger tools available on the 10x Genomics cloud. The uploaded FASTQ files comprising R1 and R2 with same Flowcell ID were applied with the library type “Single cell 3’ Gene Expression”. A new analysis was created using the “Cell Ranger Count v7.0.0” pipeline. The FASTQ files were aligned to the mouse transcriptome using the reference “Mouse (mm10) 2020-A” database with Cell Ranger count. In order to compare the cell clusters between untreated (Ctrl), *Bb*LP treated at 1 hr (LP1) and 4 hrs (LP4) their corresponding Cell ranger counts from each sample were aggregated together using “Cell Ranger Aggr v7.0.0” pipeline. Further, the CLOUPE files were downloaded, and the data analysis and visualization were carried out using Loupe cell browser v6.1.0. All raw data files are deposited in NIH GEO under the accession number GSE253285.

### scRNA-Seq Data Analysis using Seurat

Barcodes and matrix files from each sample were processed and integrated with Seurat (v5.1.0) and dplyr (v1.1.4) using R (v4.4.1) in RStudio (2024.04.2). UMAPs integrated with Harmony (v1.2.1) were created to visualize the gene expression patterns of selected genes. Heatmaps were generated using the pheatmap package (v1.0.12), pseudotime plots were created with the monocle3 package (v1.3.7), and various plots were produced using ggplot2 (v3.5.1).

### Proteomics analysis

To compare the proteomics profile with the single cell transcriptomes, 1x10^6^ BMDM were treated with 1µg/mL of *Bb*LP or PBS (for 1 or 4 hrs) in a 6-well plate. Supernatant was removed and cells were washed twice with PBS ensuring extracellular *Bb*LP (either not bound or internalized by BMDMs) are excluded from the lysates used for proteomic analysis. Total protein lysates were prepared using RIPA (Radio-Immunoprecipitation Assay) Buffer (Pierce, Thermo Fisher) supplemented with protease inhibitor (Pierce, Thermo Fisher) and stored at -20°C. Total proteins profiles and the integrity of proteins were analyzed using 5 µL of lysates on SDS-PAGE gel. The samples were submitted to proteomics core at UTMB for mass spectrometry analysis. The protein sample digestion, LC/MS analysis and data analysis were performed using previously established methods [147], with slight modifications as follows:

### Protein sample digestion

Each protein sample mixture was solubilized with 5% SDS, 50 mM Triethylammonium bicarbonate (TEAB), pH 7.55, final volume 25 μl. The sample was then centrifuged at 17,000 g for 10 min to remove any debris. Proteins were reduced by treatment with 20 mM Tris(2-carboxyethyl)phosphine hydrochloride (TCEP-HCL, Thermo Scientific) and incubated at 65 °C for 30 min. The sample was cooled to room temperature and 1 μl of 0.5 M iodoacetamide acid added and allowed to react for 20 min in the dark. 2.75 μl of 12% phosphoric acid was added to the protein solution. 165 μl of binding buffer (90% Methanol, 100 mM TEAB; pH 7.1) was then added to the solution. The resulting solution was added to S-Trap spin column (protifi.com) and passed through the column by centrifugation using a bench top centrifuge (30 s spin at 4000×g). The spin column was washed with 400 μl of binding buffer and centrifuged. The column was washed with binding buffer two more times. Trypsin was added to the protein mixture in a ratio of 1:25 in 50 mM TEAB, pH 8, and incubated at 37 °C for 4 h. Peptides were eluted with 75 μl of 50% acetonitrile, 0.2% formic acid, and then washed again with 75 μl of 80% acetonitrile, 0.2% formic acid. The combined peptide solution was then dried in a speed vac and resuspended in 2% acetonitrile, 0.1% formic acid, 97.9% water and placed in an autosampler vial for Mass Spec analysis.

### NanoLC MS/MS Analysis

Peptide mixtures were analyzed by nanoflow liquid chromatography-tandem mass spectrometry (nanoLC-MS/MS) using an UltiMate 3000 RSLCnano system (Dionex), coupled to a Thermo Orbitrap Fusion mass spectrometer (Thermo Fisher Scientific, San Jose, CA) via a nanospray ion source. A trap-and-elute method was employed with a C18 PepMap100 trap column (300 µm x 5 mm, 5 µm particle size, Thermo Scientific) and an Acclaim PepMap 100 analytical column (75 µm x 25 cm, Thermo Scientific). After equilibrating the column with 98% solvent A (0.1% formic acid in water) and 2% solvent B (0.1% formic acid in acetonitrile), 1 µL of sample in solvent A was injected onto the trap column. Elution onto the C18 column occurred at a flow rate of 300 nL/min under the following gradient conditions: isocratic at 2% B (0-5 min); 2% to 4% B (5-6 min); 4% to 32% B (5-120 min); 32% to 90% B (120-123 min); isocratic at 90% B (123-126 min); 90% to 2% B (126-129 min); isocratic at 2% B (129-130 min); 2% to 90% B (130-134 min); isocratic at 90% B (134-137 min); 90% to 2% B (137-140 min); and isocratic at 2% B (140-145 min).

LC-MS/MS data were acquired using Xcalibur (version 4.4.16.14, Thermo Scientific) in positive ion mode with a top speed data-dependent acquisition (DDA) method and a 3-second cycle time. Survey scans (m/z 375-1500) were performed in the Orbitrap at 120,000 resolution (at m/z = 400) in profile mode, with automatic maximum injection time and a normalized AGC target of 100%. The S-lens RF level was set to 60. Precursors were isolated in the quadrupole using a 1.6 Da window, and CID MS/MS was performed in centroid mode, detected in the ion trap (Rapid scan rate), with 30% CID collision energy. Maximum injection time was set to Dynamic with a normalized AGC target of 20%. Monoisotopic precursor selection (MIPS) and charge state filtering (charge states 2-7) were enabled. Dynamic exclusion was applied to exclude selected precursor ions within a ±10 ppm mass tolerance for 30 seconds after MS/MS acquisition.

### Proteomic Data Analysis

Tandem mass spectra were extracted and charge state deconvolved using Proteome Discoverer (version 2.5.0.402, Thermo Fisher). All MS/MS spectra were searched against a concatenated FASTA database comprising the Uniprot mouse database (version 03-29-2016) and the Uniprot *B. burgdorferi* database (downloaded 09-16-2022) using Sequest. The search parameters included a parent ion tolerance of 10 ppm and a fragment ion tolerance of 0.60 Da. Trypsin was specified as the enzyme, allowing up to two missed cleavages. Carbamidomethylation of cysteine was set as a fixed modification, while oxidation of methionine and deamidation of asparagine and glutamine were set as variable modifications. Peptide identifications were filtered using Percolator to achieve a false discovery rate (FDR) of 1%. Label-free quantitation was carried out with Minora, and differential expression analysis between sample groups was performed using the Precursor Ions Quantifier node in Proteome Discoverer. Protein abundance ratios were calculated using a pairwise ratio scheme, with hypothesis testing performed via t-tests on the background population of proteins and peptides.

### Protein network analysis

To understand the key pathways that are differentially activated between untreated, LP1 and LP4 samples and to analyze the interactions between the selected DEGs, protein-protein interaction analysis and the functional enrichments were performed using the STRING database (https://string-db.org/). Briefly, the identified DEGs were submitted to the STRING database via the Multiple Proteins option and employed the mouse protein-protein interaction database from STRING online database version 11.5, and the interaction prediction was determined with high confidence (0.700). Additionally, functional ontology analysis was performed, and the key molecular functions that are found in common within the selected genes were determined. Further, the interactions between the selected genes involved in specific pathways were also predicted.

### RNA isolation and qRT-PCR analysis

RNA was isolated using RNeasy kit (Qiagen, Germany) according to the manufacturer’s protocol from BMDMs treated with or without *Bb*LP seeded in 6-well plates. One microgram of RNA per sample was reverse transcribed into cDNA using Advanced iScript cDNA synthesis kit (BioRad). cDNA was then amplified using murine gene-specific primers (Supplementary Table) and the primers for Actb gene were used as normalization control. Real-time quantitative PCR (Polymerase Chain Reaction) was performed by StepOne Plus Real-time PCR system (Applied Biosystems) using default PCR program. Results are represented as fold difference using 2^-ddct^ formula comparing the internal and the untreated controls. All results were obtained from two independent experiments with three technical replicates.

### ELISA

Levels of various cytokines such as CXCL1, CXCL2, CCL5 and IL1B present in the culture supernatants of BMDMs treated with or without *Bb*LP were determined using Mouse 1) CXCL1/KC, 2) CXCL2/MIP-2, 3) CCL5/RANTES, 4) IL1B/IL1F2 DuoSet ELISA kits (R&D Systems, Minneapolis, MN, USA), following manufacturer’s protocols. Samples were diluted accordingly to measure the OD450/570 within the linear range using Spark 10M (Tecan) multimode plate reader and recorded using Spark software.

### Immunoblot Analysis

Immunoblot analysis was performed to confirm the upregulation of DUSP1 and PD-L1 in BMDMs following treatment with borrelial lipoproteins. Briefly, 1x10^7^ BMDM cells treated with *Bb*LP for 1, 2, 4, 24 and 48 hpt. Culture supernatants were removed, and the adherent cells were washed with PBS before being lysed with RIPA buffer. The total protein concentration was quantified using Pierce BCA assay kit (ThermoFisher, USA). Equal amounts of protein (50 µg/sample) were separated using 4 to 20% polyacrylamide gels and then transferred to a polyvinylidene difluoride (PVDF) membrane, blocked with blocking buffer (10% non-fat dry milk in Tris buffer (pH 7.5) containing 200 mM Tris, 1.38 M NaCl, and 0.1% Tween 20) overnight and then incubated with a primary antibody specific for DUSP1, PD-L1 (Rabbit monoclonal IgG antibodies) or ß-actin (mouse monoclonal IgG antibody) at room temperature for 2 hrs in blocking buffer and treated with horseradish peroxidase (HRP) conjugated Goat anti-Rabbit IgG secondary antibody (Cell Signaling) or Goat anti-mouse IgG secondary antibody respectively for 1 h at room temperature. The blots were developed using an Enhanced Chemiluminescence System reagents and followed by exposure to X-ray film.

### PD-L1 expression analysis by flow cytometry

The cell surface expression of *Cd274* gene encoded PD-L1 protein in bone marrow-derived macrophages (BMDMs) treated or untreated with *Bb*LP was assessed by flow cytometry. Briefly, 1 × 10^6^ BMDM cells were plated in each well of a 6-well plate and incubated overnight at 37°C with 5% CO_2_. The BMDMs were pre-treated with 10 µM BCI inhibitor for 30 minutes, followed by treatment with 0.1 µg/mL *Bb*LP. At 24 hours post-infection (hpi), the cells were washed with PBS and detached using Accutase. The cells were then stained with anti-PD-L1-PE (Cat#12-5982-83, eBioscience) for 30 minutes, washed with PBS, and resuspended in sheath fluid. Flow cytometry analysis was performed using a BD-LSRII flow cytometer following standard protocols at the UTSA Cell Analysis Core. Data processing was conducted with FlowJo software. The difference in PD-L1 cell surface expression was quantified by gating PE^hi^ events and represented as stagger offset format of histogram overlays and dot plots showing the PD-L1^hi^ population [91]. Bar graphs represent the percentage of PD-L1^hi^ population determined from three independent experiments, with error bars indicating the standard error of the mean (SEM).

### Mitochondrial ROS detection by mitoSOX assay using flow cytometry

Mitochondrial reactive oxygen species (mROS) were measured using the MitoSOX™ Red indicator (Invitrogen). MitoSOX™ Red is specifically targeted to mitochondria in live cells and is oxidized by superoxide reactive oxygen species. To assess mROS levels in macrophages, *Bb*LP-treated or untreated PMA-differentiated THP1 cells were stained with 10 μM MitoSOX™ Red according to the manufacturer’s protocol. Flow cytometry was conducted on a BD-LSRII flow cytometer. Data analysis was performed using FlowJo software, and the results were presented as median fluorescence intensity in a bar graph format. Error bars represent the standard error of the mean (SEM) across replicates.

### Immunofluorescence staining and confocal imaging

Cell surface expression of PD-L1, CD11b and intracellular levels of DUSP1 in BMDMs with or without *Bb*LP treatment was visualized by confocal imaging [91]. Briefly, 1x10^6^ BMDM cells were plated in each well on a 6-well plate pre-loaded with a coverslip, incubated overnight at 37°C under 5% CO_2_, further treated with 10 µM BCI inhibitor for 30 minutes, followed by treatment with *Bb*LP (0.1 µg/mL) [148]. BMDMs with no inhibitor or *Bb*LP treatments were maintained as appropriate controls. For PD-L1 staining, cells were incubated with *Bb*LP for 24 hours. For staining, cover slips were washed with PBS, fixed with 4% paraformaldehyde for 15 mins, blocked with 10% normal donkey serum for 30 minutes, then stained with antibody anti-CD11b-FITC, anti-PD-L1-PE for 1 hr at room temperature [149]. For DUSP1 staining, BMDM were incubated with *Bb*LP for 1 and 4 hrs, fixed with 4% PFA and permeabilized with 0.3% Triton-X 100 in PBS for 5 minutes, washed with PBS, stained with primary antibody anti-CD11b-FITC and anti-DUSP1 (Rabbit IgG) for 1 hr and counter stained with secondary antibody anti-Rabbit IgG-AF647 for 1 hr. Slides were washed with PBS to remove unbound antibodies [91]. Finally, the cells were stained with Hoechst nuclear stain for 5 minutes and washed thoroughly and mounted on a slide in ProLong™ Gold Antifade Mountant (Thermo Scientific), covered by a cover slip and allowed to cure overnight at room temperature. Microscopic analysis was performed using a Zeiss 710 NLO 2P system, and images were analyzed using Zen software.

### Statistical analysis

Graphs were prepared and statistical analysis was performed using GraphPad Prism 7.0. Statistical differences between groups were reported to be significant when the P value was less than or equal to 0.05. Data are presented as mean ± standard error of mean (SEM). All statistical analysis methods are indicated either in the figure legends or in the results section.

## Abbreviations

*Bb*LP: *Borrelia burgdorferi* Lipoproteins
BMDMs: bone marrow derived macrophages
LP1: Lipoprotein treated BMDMs for 1 hr
LP4: Lipoprotein treated BMDMs for 4 hrs
DUSP1: Dual specificity phosphatase 1
BCI: 2-benzylidene-3-(cyclohexylamino)-1-indanone hydrochloride

## Acknowledgements

This study was supported by Public Health Service Grants 5R01AI152233 (JS) and R21AI149263 (JS) from the National Institutes of Allergy and Infectious Diseases. Single-cell library preparation and sequencing was performed through the UTSA Genomics Core which is supported by NIH grant G12-MD007591, NSF grant DBI-1337513 (BH) and UTSA. The authors acknowledge the Cell Analysis Core Facility at UTSA and Dr. Sandra M. Cardona (Cell Analysis Core Director) for technical support. We would like to thank Dr. William Russell for conducting the Mass spectrometry and the data analysis at Mass Spectrometry Facility, University of Texas Medical Branch, Galveston, Texas, supported by the funding from Cancer Prevention Research Institute of Texas (Grant number RP190682). We thank Nathan Kilgore and Jolie Starling for the critical reading of the manuscript. The content is solely the responsibility of the authors and does not necessarily represent the official views of the National Institutes of Health.

## References

1. Kugeler, K.J., et al., Estimating the Frequency of Lyme Disease Diagnoses, United States, 2010-2018. Emerg Infect Dis, 2021. 27(2): p. 616-619.

2. Burgdorfer, W., et al., Lyme disease-a tick-borne spirochetosis? Science, 1982. 216(4552): p. 1317-9.

3. Radolf, J.D., et al., Of ticks, mice and men: understanding the dual-host lifestyle of Lyme disease spirochaetes. Nat Rev Microbiol, 2012. 10(2): p. 87–99.

4. Radolf, J.D., et al., Lyme Disease in Humans. Curr Issues Mol Biol, 2021. 42: p. 333–384.

5. Casselli, T., et al., A murine model of Lyme disease demonstrates that Borrelia burgdorferi colonizes the dura mater and induces inflammation in the central nervous system. PLoS Pathog, 2021. 17(2): p. e1009256.

6. Wormser, G.P., et al., The clinical assessment, treatment, and prevention of lyme disease, human granulocytic anaplasmosis, and babesiosis: clinical practice guidelines by the Infectious Diseases Society of America. Clin Infect Dis, 2006. 43(9): p. 1089–134.

7. Steere, A.C., et al., Lyme borreliosis. Nat Rev Dis Primers, 2016. 2: p. 16090.

8. Verschoor, Y.L., et al., Persistent Borrelia burgdorferi Sensu Lato Infection after Antibiotic Treatment: Systematic Overview and Appraisal of the Current Evidence from Experimental Animal Models. Clin Microbiol Rev, 2022. 35(4): p. e0007422.

9. Aucott, J.N., Posttreatment Lyme disease syndrome. Infect Dis Clin North Am, 2015. 29(2): p. 309–23.

10. Bockenstedt, L.K., et al., Spirochete antigens persist near cartilage after murine Lyme borreliosis therapy. J Clin Invest, 2012. 122(7): p. 2652–60.

11. Jutras, B.L., et al., Borrelia burgdorferi peptidoglycan is a persistent antigen in patients with Lyme arthritis. Proc Natl Acad Sci U S A, 2019. 116(27): p. 13498–13507.

12. Holub, M.N., et al., Peptidoglycan in osteoarthritis synovial tissue is associated with joint inflammation. Arthritis Research & Therapy, 2024. 26(1): p. 77.

13. Radolf, J.D., et al., Characterization of outer membranes isolated from Borrelia burgdorferi, the Lyme disease spirochete. Infect Immun, 1995. 63(6): p. 2154–63.

14. Fraser, C.M., et al., Genomic sequence of a Lyme disease spirochaete, Borrelia burgdorferi. Nature, 1997. 390(6660): p. 580-6.

15. Casjens, S., et al., A bacterial genome in flux: the twelve linear and nine circular extrachromosomal DNAs in an infectious isolate of the Lyme disease spirochete Borrelia burgdorferi. Mol Microbiol, 2000. 35(3): p. 490–516.

16. Zuckert, W.R., A call to order at the spirochaetal host-pathogen interface. Mol Microbiol, 2013. 89(2): p. 207–11.

17. Ben-Menachem, G., et al., A newly discovered cholesteryl galactoside from Borrelia burgdorferi. Proc Natl Acad Sci U S A, 2003. 100(13): p. 7913–8.

18. Hossain, H., et al., Structural analysis of glycolipids from Borrelia burgdorferi. Biochimie, 2001. 83(7): p. 683–92.

19. Kinjo, Y., et al., Natural killer T cells recognize diacylglycerol antigens from pathogenic bacteria. Nat Immunol, 2006. 7(9): p. 978–86.

20. DeHart, T.G., et al., The unusual cell wall of the Lyme disease spirochaete Borrelia burgdorferi is shaped by a tick sugar. Nat Microbiol, 2021. 6(12): p. 1583–1592.

21. Rosa, P.A., K. Tilly, and P.E. Stewart, The burgeoning molecular genetics of the Lyme disease spirochaete. Nat Rev Microbiol, 2005. 3(2): p. 129–43.

22. Gaber, A.M., et al., Comparative transcriptome analysis of Peromyscus leucopus and C3H mice infected with the Lyme disease pathogen. Front Cell Infect Microbiol, 2023. 13: p. 1115350.

23. Stevenson, B. and C.A. Brissette, Erp and Rev Adhesins of the Lyme Disease Spirochete’s Ubiquitous cp32 Prophages Assist the Bacterium during Vertebrate Infection. Infect Immun, 2023. 91(3): p. e0025022.

24. Kovryha, N., et al., Prevalence of Borrelia burgdorferi and Anaplasma phagocytophilum in Ixodid Ticks from Southeastern Ukraine. Vector Borne Zoonotic Dis, 2021. 21(4): p. 242–246.

25. Kurokawa, C., et al., Interactions between Borrelia burgdorferi and ticks. Nat Rev Microbiol, 2020. 18(10): p. 587–600.

26. Stevenson, B. and J. Seshu, Regulation of Gene and Protein Expression in the Lyme Disease Spirochete. Curr Top Microbiol Immunol, 2018. 415: p. 83–112.

27. Hyde, J.A., Borrelia burgdorferi Keeps Moving and Carries on: A Review of Borrelial Dissemination and Invasion. Front Immunol, 2017. 8: p. 114.

28. Dowdell, A.S., et al., Comprehensive Spatial Analysis of the Borrelia burgdorferi Lipoproteome Reveals a Compartmentalization Bias toward the Bacterial Surface. J Bacteriol, 2017. 199(6).

29. Pal, U., et al., OspC facilitates Borrelia burgdorferi invasion of Ixodes scapularis salivary glands. J Clin Invest, 2004. 113(2): p. 220–30.

30. Yang, X.F., et al., Essential role for OspA/B in the life cycle of the Lyme disease spirochete. J Exp Med, 2004. 199(5): p. 641–8.

31. Seshu, J., et al., Inactivation of the fibronectin-binding adhesin gene bbk32 significantly attenuates the infectivity potential of Borrelia burgdorferi. Mol Microbiol, 2006. 59(5): p. 1591–601.

32. Brissette, C.A., et al., Borrelia burgdorferi RevA antigen binds host fibronectin. Infect Immun, 2009. 77(7): p. 2802–12.

33. Fischer, J.R., et al., Decorin-binding proteins A and B confer distinct mammalian cell type-specific attachment by Borrelia burgdorferi, the Lyme disease spirochete. Proc Natl Acad Sci U S A, 2003. 100(12): p. 7307–12.

34. Wu, J., et al., Invasion of eukaryotic cells by Borrelia burgdorferi requires beta(1) integrins and Src kinase activity. Infect Immun, 2011. 79(3): p. 1338–48.

35. Setubal, J.C., et al., Lipoprotein computational prediction in spirochaetal genomes. Microbiology (Reading), 2006. 152(Pt 1): p. 113–121.

36. Haake, D.A. and W.R. Zuckert, Spirochetal Lipoproteins in Pathogenesis and Immunity. Curr Top Microbiol Immunol, 2018. 415: p. 239–271.

37. Bockenstedt, L.K., R.M. Wooten, and N. Baumgarth, Immune Response to Borrelia: Lessons from Lyme Disease Spirochetes. Curr Issues Mol Biol, 2021. 42: p. 145–190.

38. Schroder, N.W., et al., Immune responses induced by spirochetal outer membrane lipoproteins and glycolipids. Immunobiology, 2008. 213(3-4): p. 329–40.

39. Benjamin, S.J., et al., Macrophage mediated recognition and clearance of Borrelia burgdorferi elicits MyD88-dependent and -independent phagosomal signals that contribute to phagocytosis and inflammation. BMC Immunology, 2021. 22(1): p. 32.

40. Salazar, J.C., et al., Activation of human monocytes by live Borrelia burgdorferi generates TLR2-dependent and -independent responses which include induction of IFN-beta. PLoS Pathog, 2009. 5(5): p. e1000444.

41. Moore, M.W., et al., Phagocytosis of Borrelia burgdorferi and Treponema pallidum potentiates innate immune activation and induces gamma interferon production. Infect Immun, 2007. 75(4): p. 2046–62.

42. Cervantes, J.L., et al., Phagosomal signaling by Borrelia burgdorferi in human monocytes involves Toll-like receptor (TLR) 2 and TLR8 cooperativity and TLR8-mediated induction of IFN-beta. Proc Natl Acad Sci U S A, 2011. 108(9): p. 3683–8.

43. Farris, L.C., et al., Borrelia burgdorferi Engages Mammalian Type I IFN Responses via the cGAS-STING Pathway. J Immunol, 2023. 210(11): p. 1761–1770.

44. Kawai, T. and S. Akira, The role of pattern-recognition receptors in innate immunity: update on Toll-like receptors. Nature Immunology, 2010. 11(5): p. 373–384.

45. Shin, O.S., et al., Distinct roles for MyD88 and Toll-like receptors 2, 5, and 9 in phagocytosis of Borrelia burgdorferi and cytokine induction. Infect Immun, 2008. 76(6): p. 2341-51.

46. Miller, J.C., et al., Gene expression profiling provides insights into the pathways involved in inflammatory arthritis development: murine model of Lyme disease. Exp Mol Pathol, 2008. 85(1): p. 20–7.

47. Miller, J.C., et al., The Lyme disease spirochete Borrelia burgdorferi utilizes multiple ligands, including RNA, for interferon regulatory factor 3-dependent induction of type I interferon-responsive genes. Infect Immun, 2010. 78(7): p. 3144–53.

48. Hastey, C.J., et al., MyD88- and TRIF-independent induction of type I interferon drives naive B cell accumulation but not loss of lymph node architecture in Lyme disease. Infect Immun, 2014. 82(4): p. 1548–58.

49. Woitzik, P. and S. Linder, Molecular Mechanisms of Borrelia burgdorferi Phagocytosis and Intracellular Processing by Human Macrophages. Biology (Basel), 2021. 10(7).

50. Lasky, C.E., R.M. Olson, and C.R. Brown, Macrophage Polarization during Murine Lyme Borreliosis. Infect Immun, 2015. 83(7): p. 2627–35.

51. Montgomery, R.R., et al., Recruitment of macrophages and polymorphonuclear leukocytes in Lyme carditis. Infect Immun, 2007. 75(2): p. 613–20.

52. Greenmyer, J.R., et al., Primary Human Microglia Are Phagocytically Active and Respond to Borrelia burgdorferi With Upregulation of Chemokines and Cytokines. Front Microbiol, 2018. 9: p. 811.

53. Papalexi, E. and R. Satija, Single-cell RNA sequencing to explore immune cell heterogeneity. Nat Rev Immunol, 2018. 18(1): p. 35–45.

54. Tikhonova, A.N., et al., The bone marrow microenvironment at single-cell resolution. Nature, 2019. 569(7755): p. 222-228.

55. Hermann, B.P., et al., The Mammalian Spermatogenesis Single-Cell Transcriptome, from Spermatogonial Stem Cells to Spermatids. Cell Rep, 2018. 25(6): p. 1650–1667 e8.

56. Mutoji, K., et al., TSPAN8 Expression Distinguishes Spermatogonial Stem Cells in the Prepubertal Mouse Testis. Biol Reprod, 2016. 95(6): p. 117.

57. Helble, J.D., et al., Single-cell RNA sequencing of murine ankle joints over time reveals distinct transcriptional changes following Borrelia burgdorferi infection. iScience, 2023. 26(11): p. 108217.

58. Jiang, R., et al., Single-cell immunophenotyping of the skin lesion erythema migrans identifies IgM memory B cells. JCI Insight, 2021. 6(12).

59. Kumaresan, V., et al., Cellular and transcriptome signatures unveiled by single-cell RNA-Seq following ex vivo infection of murine splenocytes with Borrelia burgdorferi. Front Immunol, 2023. 14: p. 1296580.

60. Del Dotto, V., et al., Variants in Human ATP Synthase Mitochondrial Genes: Biochemical Dysfunctions, Associated Diseases, and Therapies. Int J Mol Sci, 2024. 25(4).

61. Deng, Y., et al., Glucocorticoid receptor regulates PD-L1 and MHC-I in pancreatic cancer cells to promote immune evasion and immunotherapy resistance. Nature Communications, 2021. 12(1): p. 7041.

62. Cheng, S.W., et al., GBP5 Repression Suppresses the Metastatic Potential and PD-L1 Expression in Triple-Negative Breast Cancer. Biomedicines, 2021. 9(4).

63. Daines, M.O. and G.K. Hershey, A novel mechanism by which interferon-gamma can regulate interleukin (IL)-13 responses. Evidence for intracellular stores of IL-13 receptor alpha -2 and their rapid mobilization by interferon-gamma. J Biol Chem, 2002. 277(12): p. 10387-93.

64. Wang, J., et al., A transcriptional program associated with cell cycle regulation predominates in the anti-inflammatory effects of CX-5461 in macrophage. Front Pharmacol, 2022. 13: p. 926317.

65. Bansal, A., et al., Interplay between nuclear factor-&#x3ba;B, p38 MAPK, and glucocorticoid receptor signaling synergistically induces functional TLR2 in lung epithelial cells. Journal of Biological Chemistry, 2022. 298(4).

66. Mutlu, M., et al., Small molecule induced STING degradation facilitated by the HECT ligase HERC4. Nature Communications, 2024. 15(1): p. 4584.

67. Bolz, D.D., et al., Dual role of MyD88 in rapid clearance of relapsing fever Borrelia spp. Infect Immun, 2006. 74(12): p. 6750–60.

68. Giambartolomei, G.H., et al., Induction of pro- and anti-inflammatory cytokines by Borrelia burgdorferi lipoproteins in monocytes is mediated by CD14. Infect Immun, 1999. 67(1): p. 140–7.

69. Gupta, A., et al., A human secretome library screen reveals a role for Peptidoglycan Recognition Protein 1 in Lyme borreliosis. PLoS Pathog, 2020. 16(11): p. e1009030.

70. Chen, Y., et al., Borrelia peptidoglycan interacting Protein (BpiP) contributes to the fitness of Borrelia burgdorferi against host-derived factors and influences virulence in mouse models of Lyme disease. PLoS Pathog, 2021. 17(4): p. e1009535.

71. Guyard, C., et al., Periplasmic flagellar export apparatus protein, FliH, is involved in post-transcriptional regulation of FlaB, motility and virulence of the relapsing fever spirochete Borrelia hermsii. PLoS One, 2013. 8(8): p. e72550.

72. Pereira, M.J., et al., Lipoproteome screening of the Lyme disease agent identifies inhibitors of antibody-mediated complement killing. Proc Natl Acad Sci U S A, 2022. 119(13): p. e2117770119.

73. Booth, C.E., Jr., et al., Borrelia miyamotoi FbpA and FbpB Are Immunomodulatory Outer Surface Lipoproteins With Distinct Structures and Functions. Front Immunol, 2022. 13: p. 886733.

74. Wang, Q., et al., Fibroblast growth factor 13 stabilizes microtubules to promote Na(+) channel function in nociceptive DRG neurons and modulates inflammatory pain. J Adv Res, 2021. 31: p. 97–111.

75. Bonacina, F., et al., Myeloid apolipoprotein E controls dendritic cell antigen presentation and T cell activation. Nat Commun, 2018. 9(1): p. 3083.

76. Banday, A.R., et al., Replication-dependent histone genes are actively transcribed in differentiating and aging retinal neurons. Cell Cycle, 2014. 13(16): p. 2526–41.

77. Mould, K.J., et al., Single cell RNA sequencing identifies unique inflammatory airspace macrophage subsets. JCI Insight, 2019. 4(5).

78. Yao, W., et al., Single Cell RNA Sequencing Identifies a Unique Inflammatory Macrophage Subset as a Druggable Target for Alleviating Acute Kidney Injury. Adv Sci (Weinh), 2022. 9(12): p. e2103675.

79. Tian, Z. and S. Yang, Integrating the characteristic genes of macrophage pseudotime analysis in single-cell RNA-seq to construct a prediction model of atherosclerosis. Aging (Albany NY), 2023. 15(13): p. 6361–6379.

80. Takiguchi, H., et al., Macrophages with reduced expressions of classical M1 and M2 surface markers in human bronchoalveolar lavage fluid exhibit pro-inflammatory gene signatures. Scientific Reports, 2021. 11(1): p. 8282.

81. Wang, J., et al., HMGB1 participates in LPS-induced acute lung injury by activating the AIM2 inflammasome in macrophages and inducing polarization of M1 macrophages via TLR2, TLR4, and RAGE/NF-κB signaling pathways. Int J Mol Med, 2020. 45(1): p. 61–80.

82. Pérez, S. and S. Rius-Pérez, Macrophage Polarization and Reprogramming in Acute Inflammation: A Redox Perspective. Antioxidants (Basel), 2022. 11(7).

83. Carreras-Gonzalez, A., et al., Regulation of macrophage activity by surface receptors contained within Borrelia burgdorferi-enriched phagosomal fractions. PLoS Pathog, 2019. 15(11): p. e1008163.

84. Barriales, D., et al., Borrelia burgdorferi infection induces long-term memory-like responses in macrophages with tissue-wide consequences in the heart. PLoS Biol, 2021. 19(1): p. e3001062.

85. Tullai, J.W., et al., Identification of transcription factor binding sites upstream of human genes regulated by the phosphatidylinositol 3-kinase and MEK/ERK signaling pathways. J Biol Chem, 2004. 279(19): p. 20167–77.

86. Nguyen, H.Q., B. Hoffman-Liebermann, and D.A. Liebermann, The zinc finger transcription factor Egr-1 is essential for and restricts differentiation along the macrophage lineage. Cell, 1993. 72(2): p. 197–209.

87. Khachigian, L.M., Early Growth Response-1, an Integrative Sensor in Cardiovascular and Inflammatory Disease. Journal of the American Heart Association, 2021. 10(22): p. e023539.

88. Wang, L., et al., An essential role of PI3K in the control of West Nile virus infection. Scientific Reports, 2017. 7(1): p. 3724.

89. Lochhead, R.B., et al., Lyme arthritis: linking infection, inflammation and autoimmunity. Nature Reviews Rheumatology, 2021. 17(8): p. 449–461.

90. Yang, K., et al., Integrating systematic biological and proteomics strategies to explore the pharmacological mechanism of danshen yin modified on atherosclerosis. Journal of Cellular and Molecular Medicine, 2020. 24(23): p. 13876–13898.

91. Wang, L., et al., PD-L1-expressing tumor-associated macrophages are immunostimulatory and associate with good clinical outcome in human breast cancer. Cell Reports Medicine, 2024. 5(2).

92. Caratti, G., et al., Glucocorticoid activation of anti-inflammatory macrophages protects against insulin resistance. Nat Commun, 2023. 14(1): p. 2271.

93. Mantuano, E., et al., LDL receptor-related protein-1 regulates NFκB and microRNA-155 in macrophages to control the inflammatory response. Proceedings of the National Academy of Sciences, 2016. 113(5): p. 1369–1374.

94. Gandhi, H., et al., Dynamics and interaction of interleukin-4 receptor subunits in living cells. Biophys J, 2014. 107(11): p. 2515–27.

95. Umeshita-Suyama, R., et al., Characterization of IL-4 and IL-13 signals dependent on the human IL-13 receptor alpha chain 1: redundancy of requirement of tyrosine residue for STAT3 activation. Int Immunol, 2000. 12(11): p. 1499–509.

96. Petzke, M.M., et al., Borrelia burgdorferi induces a type I interferon response during early stages of disseminated infection in mice. BMC Microbiol, 2016. 16: p. 29.

97. Hoffmann, A., et al., The IkappaB-NF-kappaB signaling module: temporal control and selective gene activation. Science, 2002. 298(5596): p. 1241-5.

98. Zhao, T., et al., The NEMO adaptor bridges the nuclear factor-kappaB and interferon regulatory factor signaling pathways. Nat Immunol, 2007. 8(6): p. 592–600.

99. Kenedy, M.R., T.R. Lenhart, and D.R. Akins, The role of Borrelia burgdorferi outer surface proteins. FEMS Immunol Med Microbiol, 2012. 66(1): p. 1–19.

100. Thompson, D., J.A. Watt, and C.A. Brissette, Host transcriptome response to Borrelia burgdorferi sensu lato. Ticks Tick Borne Dis, 2021. 12(2): p. 101638.

101. Seternes, O.M., A.M. Kidger, and S.M. Keyse, Dual-specificity MAP kinase phosphatases in health and disease. Biochim Biophys Acta Mol Cell Res, 2019. 1866(1): p. 124–143.

102. Hammer, M., et al., Dual specificity phosphatase 1 (DUSP1) regulates a subset of LPS-induced genes and protects mice from lethal endotoxin shock. J Exp Med, 2006. 203(1): p. 15–20.

103. Molina, G., et al., Zebrafish chemical screening reveals an inhibitor of Dusp6 that expands cardiac cell lineages. Nat Chem Biol, 2009. 5(9): p. 680–7.

104. Oosting, M., et al., Borrelia species induce inflammasome activation and IL-17 production through a caspase-1-dependent mechanism. European Journal of Immunology, 2011. 41(1): p. 172–181.

105. Oosting, M., et al., Murine Borrelia arthritis is highly dependent on ASC and caspase-1, but independent of NLRP3. Arthritis Res Ther, 2012. 14(6): p. R247.

106. Smallie, T., et al., Dual-Specificity Phosphatase 1 and Tristetraprolin Cooperate To Regulate Macrophage Responses to Lipopolysaccharide. The Journal of Immunology, 2015. 195(1): p. 277–288.

107. Acosta, J.C., et al., Chemokine signaling via the CXCR2 receptor reinforces senescence. Cell, 2008. 133(6): p. 1006–18.

108. Chien, Y., et al., Control of the senescence-associated secretory phenotype by NF-κB promotes senescence and enhances chemosensitivity. Genes Dev, 2011. 25(20): p. 2125–36.

109. Kang, T.W., et al., Senescence surveillance of pre-malignant hepatocytes limits liver cancer development. Nature, 2011. 479(7374): p. 547-51.

110. Nanda, S.K., et al., IKKβ is required for the formation of the NLRP3 inflammasome. EMBO Rep, 2021. 22(10): p. e50743.

111. Bawazeer, M.A. and T.C. Theoharides, IL-33 stimulates human mast cell release of CCL5 and CCL2 via MAPK and NF-κB, inhibited by methoxyluteolin. European Journal of Pharmacology, 2019. 865: p. 172760.

112. Lim, S., et al., TRAF6 mediates IL-1β/LPS-induced suppression of TGF-β signaling through its interaction with the type III TGF-β receptor. PLoS One, 2012. 7(3): p. e32705.

113. Patterson, B.K., et al., Disruption of the CCL5/RANTES-CCR5 Pathway Restores Immune Homeostasis and Reduces Plasma Viral Load in Critical COVID-19. medRxiv, 2020.

114. Dinarello, C.A., A. Simon, and J.W. van der Meer, Treating inflammation by blocking interleukin-1 in a broad spectrum of diseases. Nat Rev Drug Discov, 2012. 11(8): p. 633–52.

115. Zhou, Z., et al., Single-cell profiling identifies IL1Bhi macrophages associated with inflammation in PD-1 inhibitor-induced inflammatory arthritis. Nature Communications, 2024. 15(1): p. 2107.

116. Miller, J.C., et al., A Critical Role for Type I IFN in Arthritis Development following Borrelia burgdorferi Infection of Mice1. The Journal of Immunology, 2008. 181(12): p. 8492–8503.

117. Marques, A., et al., Transcriptome Assessment of Erythema Migrans Skin Lesions in Patients With Early Lyme Disease Reveals Predominant Interferon Signaling. The Journal of Infectious Diseases, 2017. 217(1): p. 158–167.

118. Tretina, K., et al., Interferon-induced guanylate-binding proteins: Guardians of host defense in health and disease. J Exp Med, 2019. 216(3): p. 482–500.

119. Braun, E., et al., Guanylate-Binding Proteins 2 and 5 Exert Broad Antiviral Activity by Inhibiting Furin-Mediated Processing of Viral Envelope Proteins. Cell Rep, 2019. 27(7): p. 2092–2104.e10.

120. Shenoy, A.R., et al., GBP5 promotes NLRP3 inflammasome assembly and immunity in mammals. Science, 2012. 336(6080): p. 481-5.

121. Helble, J.D., et al., The PD-1/PD-L1 pathway is induced during Borrelia burgdorferi infection and inhibits T cell joint infiltration without compromising bacterial clearance. PLoS Pathog, 2022. 18(10): p. e1010903.

122. Barley, T.J., et al., Mitogen-activated protein kinase phosphatase-1 controls PD-L1 expression by regulating type I interferon during systemic Escherichia coli infection. J Biol Chem, 2022. 298(5): p. 101938.

123. Luo, Q., et al., Elevated expression of PD-1 on T cells correlates with disease activity in rheumatoid arthritis. Mol Med Rep, 2018. 17(2): p. 3297–3305.

124. Zhang, F., et al., DUSP6 Inhibitor (E/Z)-BCI Hydrochloride Attenuates Lipopolysaccharide-Induced Inflammatory Responses in Murine Macrophage Cells via Activating the Nrf2 Signaling Axis and Inhibiting the NF-κB Pathway. Inflammation, 2019. 42(2): p. 672–681.

125. Liu, Z., et al., DUSP1 mediates BCG induced apoptosis and inflammatory response in THP-1 cells via MAPKs/NF-κB signaling pathway. Scientific Reports, 2023. 13(1): p. 2606.

126. Arumugam, P., et al., The mitochondrial gene-CMPK2 functions as a rheostat for macrophage homeostasis. Front Immunol, 2022. 13: p. 935710.

127. Lai, J.H., et al., Mitochondrial CMPK2 mediates immunomodulatory and antiviral activities through IFN-dependent and IFN-independent pathways. iScience, 2021. 24(6): p. 102498.

128. Kim, H., et al., UMP-CMP kinase 2 gene expression in macrophages is dependent on the IRF3-IFNAR signaling axis. PLoS One, 2021. 16(10): p. e0258989.

129. Tokarz, P., et al., PARP1-LSD1 functional interplay controls transcription of SOD2 that protects human pro-inflammatory macrophages from death under an oxidative condition. Free Radical Biology and Medicine, 2019. 131: p. 218–224.

130. Kerstholt, M., et al., Borrelia burgdorferi inhibits NADPH-mediated reactive oxygen species production through the mTOR pathway. Ticks and Tick-borne Diseases, 2022. 13(4): p. 101943.

131. Wawrzeniak, K., et al., Effect of Borrelia burgdorferi Outer Membrane Vesicles on Host Oxidative Stress Response. Antibiotics (Basel), 2020. 9(5).

132. Kinugasa, H., et al., Mitochondrial SOD2 regulates epithelial–mesenchymal transition and cell populations defined by differential CD44 expression. Oncogene, 2015. 34(41): p. 5229–5239.

133. Bauerfeld, C., et al., MKP-1 Modulates Mitochondrial Transcription Factors, Oxidative Phosphorylation, and Glycolysis. Immunohorizons, 2020. 4(5): p. 245–258.

134. Morgan, M.J. and Z.-g. Liu, Crosstalk of reactive oxygen species and NF-κB signaling. Cell Research, 2011. 21(1): p. 103–115.

135. Talwar, H., et al., MKP-1 negatively regulates LPS-mediated IL-1β production through p38 activation and HIF-1α expression. Cell Signal, 2017. 34: p. 1–10.

136. Schilperoort, M., et al., The role of efferocytosis-fueled macrophage metabolism in the resolution of inflammation. Immunological Reviews, 2023. 319(1): p. 65–80.

137. Cai, S., et al., Mitochondrial dysfunction in macrophages promotes inflammation and suppresses repair after myocardial infarction. The Journal of Clinical Investigation, 2023. 133(4).

138. Zanoni, P., et al., Endocytosis of lipoproteins. Atherosclerosis, 2018. 275: p. 273–295.

139. Raju, B.V., et al., Oligopeptide permease A5 modulates vertebrate host-specific adaptation of Borrelia burgdorferi. Infect Immun, 2011. 79(8): p. 3407–20.

140. Van Laar, T.A., et al., Effect of levels of acetate on the mevalonate pathway of Borrelia burgdorferi. PLoS One, 2012. 7(5): p. e38171.

141. Van Laar, T.A., et al., Statins reduce spirochetal burden and modulate immune responses in the C3H/HeN mouse model of Lyme disease. Microbes Infect, 2016. 18(6): p. 430–435.

142. Karna, S.L., et al., CsrA modulates levels of lipoproteins and key regulators of gene expression critical for pathogenic mechanisms of Borrelia burgdorferi. Infect Immun, 2011. 79(2): p. 732–44.

143. Karna, S.L., et al., Contributions of environmental signals and conserved residues to the functions of carbon storage regulator A of Borrelia burgdorferi. Infect Immun, 2013. 81(8): p. 2972–85.

144. Miller, C.L., S.L. Karna, and J. Seshu, Borrelia host adaptation Regulator (BadR) regulates rpoS to modulate host adaptation and virulence factors in Borrelia burgdorferi. Mol Microbiol, 2013. 88(1): p. 105–24.

145. Caine, J.A., et al., Borrelia burgdorferi outer surface protein C (OspC) binds complement component C4b and confers bloodstream survival. Cellular Microbiology, 2017. 19(12): p. e12786.

146. Goffi, F., et al., The inhibitor of I kappa B alpha phosphorylation BAY 11-7082 prevents NMDA neurotoxicity in mouse hippocampal slices. Neurosci Lett, 2005. 377(3): p. 147–51.

147. Deberneh, H.M., et al., A large-scale LC-MS dataset of murine liver proteome from time course of heavy water metabolic labeling. Scientific Data, 2023. 10(1): p. 635.

148. Benito-León, M., et al., BCI, an inhibitor of the DUSP1 and DUSP6 dual specificity phosphatases, enhances P2X7 receptor expression in neuroblastoma cells. Front Cell Dev Biol, 2022. 10: p. 1049566.

149. Lai, C.Y., et al., Different Induction of PD-L1 (CD274) and PD-1 (CD279) Expression in THP-1-Differentiated Types 1 and 2 Macrophages. J Inflamm Res, 2021. 14: p. 5241–5249.

